# Trabectedin Enhances Oncolytic Virotherapy by Reducing Barriers to Virus Spread and Cytotoxic Immunity in Preclinical Pediatric Bone Sarcoma

**DOI:** 10.1101/2024.03.02.582994

**Authors:** Emily M. Ringwalt, Mark A. Currier, Andrea M. Glaspell, Chun-Yu Chen, Matthew V. Cannon, Maren Cam, Amy C. Gross, Matthew Gust, Pin-Yi Wang, Louis Boon, Laura E. Biederman, Emily Schwarz, Prajwal Rajappa, Dean A. Lee, Elaine R. Mardis, William E. Carson, Ryan D. Roberts, Timothy P. Cripe

## Abstract

We previously reported that the DNA alkylator and transcriptional-blocking chemotherapeutic agent trabectedin enhances oncolytic herpes simplex viroimmunotherapy in human sarcoma xenograft models, though the mechanism remained to be elucidated. Here we report trabectedin disrupts the intrinsic cellular anti-viral response which increases viral transcript spread throughout the human tumor cells. We also extended our synergy findings to syngeneic murine sarcoma models, which are poorly susceptible to virus infection. In the absence of robust virus replication, we found trabectedin enhanced viroimmunotherapy efficacy by reducing immunosuppressive macrophages and stimulating granzyme expression in infiltrating T and NK cells to cause immune-mediated tumor regressions. Thus, trabectedin enhances both the direct virus-mediated killing of tumor cells and the viral-induced activation of cytotoxic effector lymphocytes to cause tumor regressions across models. Our data provide a strong rationale for clinical translation as both mechanisms should be simultaneously active in human patients.

## Introduction

Immunotherapies are transforming the treatment of cancer, especially for patients with aggressive disease. Numerous advances in immunotherapy seek to strengthen the immune recognition of cancer. Oncolytic viroimmunotherapy leverages both a direct virus-mediated killing of tumor cells and a stimulation of antitumor immunity. Oncolytic virotherapies selectively replicate within cancer cells, often by taking advantage of dysregulated signaling pathways. Currently, the only FDA-approved oncolytic virotherapy is talimogene laherparepvec (T-VEC), a modified oncolytic herpes simplex virus (oHSV) used to treat recurrent melanoma.^1,2^ New virotherapies in clinical trials take advantage of the targetability and immune-stimulating effects these therapies possess and seek to boost clinical efficacy by elucidating potential combination approaches.^3,4^

Pediatric and young adult patients with aggressive solid tumors continue to experience poor outcomes and are a population that may benefit from therapies such as oHSV. In particular, outcomes for Ewing sarcoma and osteosarcoma—the two most common pediatric tumors of the bone and surrounding soft tissue—have not improved in decades, despite collaborative international clinical trial consortia. Advanced and metastatic disease develops in 25% of patients despite standard-of-care therapy.^5–9^ Long-term survival rates for these cases stagnate around 20-40%.^5,7,10^ Osteosarcoma typically forms due to sporadic mutations, often those leading to loss of tumor suppressor function and diverse phenotypic heterogeneity that supports tumor survival and metastasis.^11,12^ This malignancy primarily metastasizes to the lungs where lung-osteosarcoma interactions support colonization and survival.^13,14^ Ewing sarcoma is characterized by a chromosomal translocation that creates a fusion oncogene of the *EWS* gene with a member of the ETS family of transcription factors, typically *FLI1.*^15^ The resulting fusion oncoprotein acts an aberrant transcription factor and chromatin remodeler.^15^

Oncolytic viroimmunotherapy, including oHSV, has exhibited preclinical efficacy against pediatric bone sarcoma.^16^ We found that HSV1716 (Seprehvir) displayed clinical safety and evidence of immune activation when utilized to treat pediatric solid tumors.^17,18^ However, durable virotherapy responses remained elusive, possibly due to a rapid virus clearance by anti-viral immune cells and the presence of immunosuppressive cells, such as tumor-associated macrophages and myeloid-derived suppressor cells (MDSC), that reside within the tumor microenvironment and are enriched by tumor signaling and persistent immune activation. Thus, while clinically safe, depleting or inhibiting the immunosuppressive myeloid cells within the tumor microenvironment may be required to achieve oHSV efficacy in pediatric sarcoma patients.

Trabectedin is an alkylating agent that displays myelolytic properties and is therefore a promising candidate to disrupt the immunosuppressive microenvironment in pediatric sarcoma patients. Trabectedin is FDA-approved for the treatment of advanced liposarcoma and leiomyosarcoma and binds the minor groove DNA to disrupt gene expression, DNA repair mechanisms and polymerase activity.^19–25^ In Ewing sarcoma, these mechanisms of action exploit vulnerabilities for oncogene dependence, leading to tumor cell death in preclinical and clinical settings.^26–30^ To achieve myelolytic effects, the drug induces apoptosis via TRAIL-R2 death receptor signaling in myeloid cells, but not in natural killer (NK) or T cells.^20,22^ Recent clinical studies found that trabectedin is safe in children and adolescents with solid tumors, with some grade 3 and 4 adverse events including fatigue, neutropenia and hepatotoxicity.^31–33^

In this study, we hypothesized that trabectedin would enhance oHSV virotherapy by reducing myeloid-derived immune suppression and increasing antitumor immune activation. Strikingly, our preclinical efficacy and survival results surpassed those we previously observed when combining myeloid depletion with oHSV, suggesting additional mechanisms underscore the combinatorial efficacy.^34^ Indeed, we define a novel mechanism for synergy whereby trabectedin enhances the direct antitumor and immune stimulatory effects of oHSV. Collectively, our data provide strong preclinical evidence supporting a combinatorial therapy consisting of oHSV and trabectedin for patients with aggressive pediatric sarcomas.

## Results

### oHSV+trabectedin therapy causes generalizable human xenograft regressions

Given the efficacy we previously observed in combining an oHSV with trabectedin against the Ewing sarcoma A673 xenograft,^34^ we sought to determine if this synergy proved generalizable to other tumor models. We observed tumor burden reductions in Ewing sarcoma xenografts (CHLA258, EW5) using the previously reported combination of the oHSV rRp450 and trabectedin (Figure 1A-B). Each tumor model responded differently to the monotherapies: the trabectedin-sensitive CHLA258 demonstrated the known susceptibility of fusion oncogene-dependent tumors to trabectedin,^26,28^ while EW5 exhibited sensitivity to oHSV alone. Despite these differences in susceptibility to monotherapy, both tumor models displayed the best overall response rates when treated with oHSV+trabectedin combination therapy, with numerous partial (PR = 19/40, 48%) and complete (CR = 7/40, 18%) responses by day 28 (Figure 1A-B, Figure S1). The disease stabilization rate (DSR = CR + PR + SD) for combination-treated EW5 (DSR = 16/19, 84%) nearly equaled that of oHSV monotherapy (DSR = 15/17, 88%), yet a higher proportion of tumors responded to the combinatorial therapy (CR + PR = 9/19, 47% vs. 6/17, 35%) (Figure S1). For CHLA258, the combination therapy conferred a significant survival advantage over the monotherapies, despite the model’s vulnerability to trabectedin alone (Figure S2). Thus, the combination of oHSV and trabectedin induced tumor regressions in these Ewing sarcoma xenograft models with varying sensitivities to each monotherapy.

**Figure 1.**
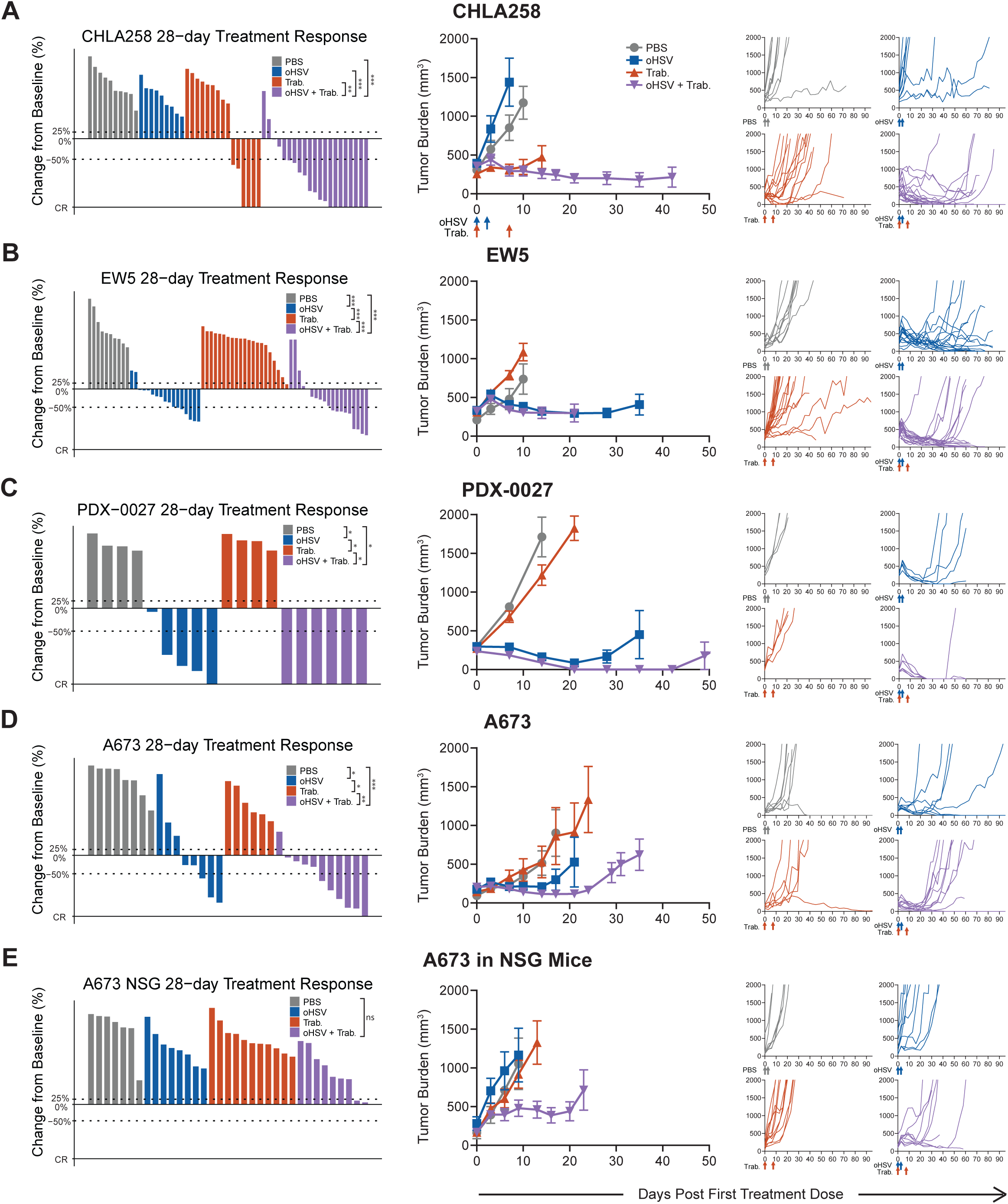
oHSV+trabectedin synergizes to increase the disease control rate and reduce tumor burden in human xenograft models. The best response for each treated tumor through 28 days, the average tumor burden, and spider plots tracking individual tumor volumes over the full study period are shown for A) CHLA-258 (Ewing sarcoma), B) EW5 (Ewing sarcoma), C) PDX-0027 (rhabdomyosarcoma), D) A673 (Ewing sarcoma), and E) A673 in NSG-SGM3 NK-deficient mice (lack T, B, and NK cells). PBS and oHSV (1.0 x 107 pfu) were given i.Tu. on Days 0 & 2. Trabectedin (0.15 mg/kg) was given I.V. on Days 0 & 7. Statistical analyses of the disease control rates (CR+PR+SD) were performed using a pairwise Fisher’s exact test with p values adjusted using the Benjamini-Hochberg procedure; *p < 0.05, **p < 0.01, ***p < 0.001. Summarized data with error bars depict mean ± SEM.

To test whether this combinatorial efficacy is maintained across oHSV constructs, we also combined trabectedin with HSV1716, an oHSV shown to be clinically safe in pediatric patients.^17^ Indeed, the combination proved efficacious against both rhabdomyosarcoma PDX-0027 (CR = 6/6, 100%) and Ewing sarcoma A673 (CR + PR = 6/11, 55%) xenografts (Figure 1C, D, Figure S1). The synergy of trabectedin with oHSV extended from numerous human xenograft models to differing oHSV constructs, indicating that trabectedin combined with oHSV offers reproducible synergistic efficacy across multiple model systems.

### Natural killer cells contribute to the combinatorial efficacy against Ewing sarcoma in immunodeficient mice

To characterize the mechanism behind the synergy between oHSV and trabectedin, we investigated the role of the immune microenvironment in the combination efficacy. If trabectedin modulated immunosuppressive myeloid cells, we anticipated that the mechanism of synergy involved an increase in activated immune effector cells. Given the nature of xenograft models, the immunodeficient (T and B cell-deficient) mouse model necessary for human tumor growth restricted this effector subset to NK cells. To determine the reliance of combination efficacy on these NK cells, we removed NK cells from the system using a mouse model deficient in NK cells (Figure 1E). In this model, combination efficacy no longer induced tumor shrinkages, though there was still evidence of survival advantage despite the absence of NK cells (Figure S2). A lower overall response rate in the absence of NK cells (SD = 2/9, 22% and CR + PR = 0/9, 0%) suggests that there is a role for NK cells in driving tumor regressions in these xenograft models (Figure S1). However, whether this mechanism operated alone or in concert with another mechanism to achieve combination efficacy required further investigation.

### Trabectedin increases oHSV spread by disrupting the intracellular intrinsic antiviral response

To gather an in-depth, high-resolution snapshot of the treatment-induced effects in the cells of the tumor microenvironment, we performed single-cell RNA sequencing (scRNAseq, n = 2) before widespread treatment-mediated cell death, which prevented analyses at later time points (Figure 2A, Figure S3). Given the known ability of trabectedin to bind to DNA and disrupt gene expression^25^, we sought to observe the effects of treatment on a transcriptional level. To capture the effects on virus gene expression, we also analyzed the treated tumors for oHSV transcript presence. A potential mechanism of synergy could be increased viral infection, yet we did not observe an increase in recoverable live virus in tumors from animals treated with the combination compared to tumors from animals treated with oHSV alone (Figure 2B). Despite the lack of increased virus titers, we found that trabectedin increased the detectable numbers of tumor cells containing oHSV and the abundance of oHSV transcripts in tumor cells (Figure 2C, D).

**Figure 2.**
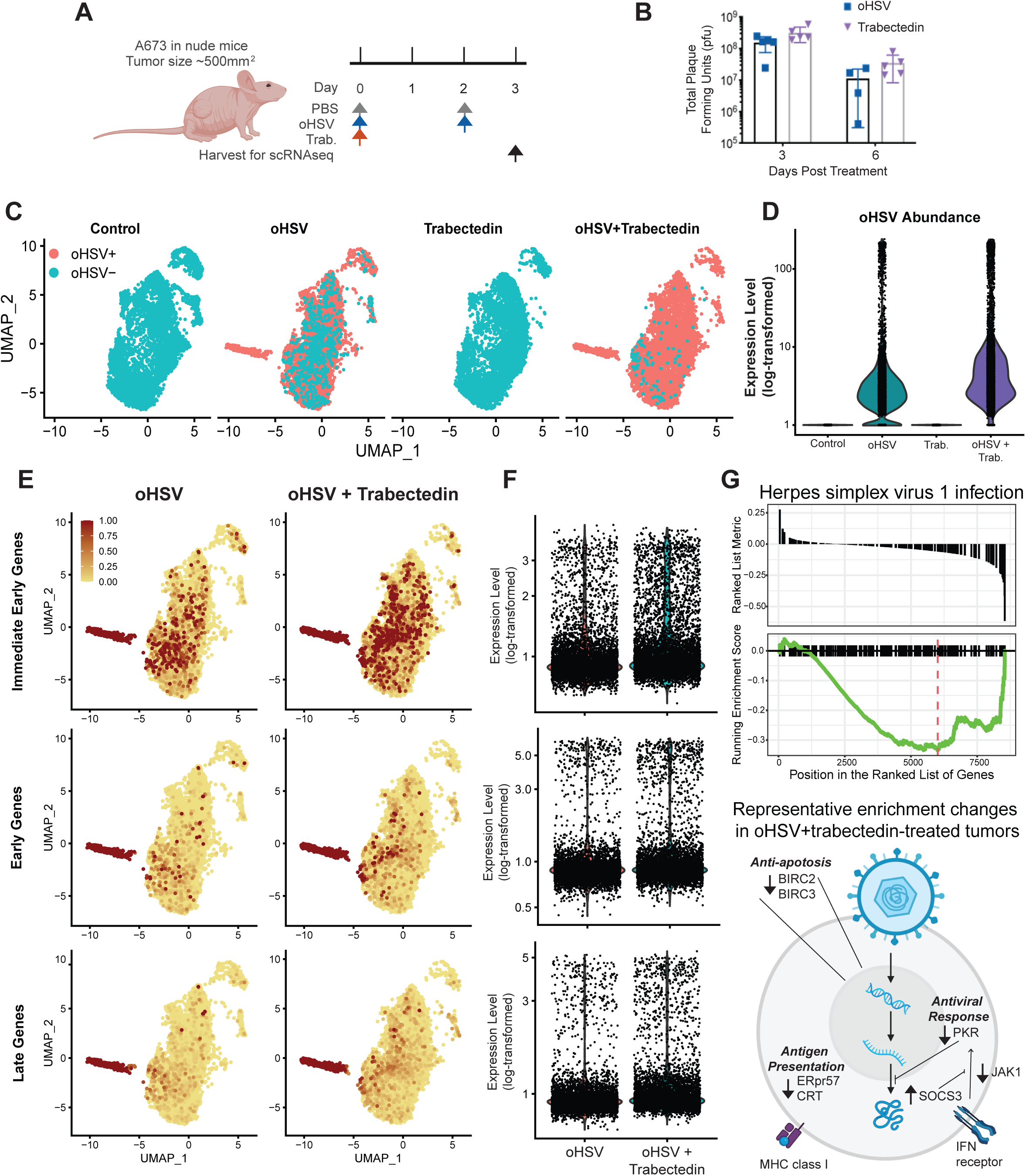
Trabectedin increases viral spread and decreases antiviral gene expression in immunodeficient Ewing sarcoma models. A) Treatment schema for scRNAseq tumor collection: PBS and oHSV (1.0 x 10^7^ pfu) were given i.Tu. on Days 0 & 2, trabectedin (0.15 mg/kg) was given I.V. on Day 0, and scRNAseq samples were collected on Day 3 to ensure sufficient cell viability for sequencing. B) oHSV viral titer (pfu) from A673 tumors treated with a single dose of oHSV or oHSV+trabectedin. C) UMAP plots of oHSV transcript presence in all tumor cells, split by treatment group. D) Violin plot of the expression level (log-transformed) of all oHSV transcripts for all tumor cells in each treatment group. E) Expression level of HSV time-dependent gene modules shown as a feature plot (UMAP) for all tumor cells and split by treatment groups that contained oHSV. A relative expression cutoff of 1 minimized color skewing by high expressing outlier cells and maintained color scale consistency between feature plots. F) Violin plot of the expression level (log-transformed) of HSV time-dependent gene modules for all tumor cells in treatment groups that contained oHSV. G) Results from gene set enrichment analysis showing the running enrichment score for the “Herpes simplex virus 1 infection” pathway, which represents the intrinsic cellular response to HSV-1. Gene set enrichment analysis was performed on a gene list which was ranked by the gene expression fold change (log2FC) calculated for tumor cells treated with oHSV+trabectedin compared to those treated with oHSV monotherapy. The schema displays the top genes with enrichment changes from the “Herpes simplex virus 1 infection” pathway.

We distinguished the three known transcriptional stages of HSV cellular infection— immediate early, early, and late genes—and calculated a module score for all the genes in each category to elucidate whether the effect of trabectedin on increased oHSV transcript spread was merely due to a leakiness of dying cells or was biologically relevant. If viral transcripts were spread in a nondiscriminatory mechanism via dying cells, we would expect cells to display a random distribution of transcripts from differing transcriptional stages of infection. In contrast, we observed that the trabectedin-mediated increase in oHSV transcript spread correlated with increases in infection stage-specific genes in a manner that reflected the timeline of cellular infection (Figure 2E, F and Figure S4). We also distinguished one cluster of highly infected and likely actively oHSV-replicating cells, as seen in the high expression of genes from all stages of oHSV infection.

To explain the trabectedin-mediated increase in oHSV transcript spread, we analyzed tumor cells for genes and pathways related to the intracellular response to the virus. We found that the combination-treated tumor cells showed a decreased enrichment of the herpes simplex virus 1 infection pathway, compared to oHSV-treated tumor cells (Fig 2G). This pathway consisted not of HSV genes, but rather intracellular responses to HSV infection. Most notably, this pathway includes intracellular antiviral responses, such as PKR signaling, and anti-apoptotic factors, both of which showed decreased expression in combination-treated tumors (Fig 2G). Trabectedin disrupted the gene expression of the antiviral response, leading to increased oHSV presence in tumor cells. Based on the lack of difference in productive viral infection between mono- and combinatorial therapies (Figure 2B), oncolysis likely occurred prior to a productive infection. Thus, the mechanism of efficacy in human Ewing sarcoma xenografts depended on trabectedin-mediated oHSV intracellular pathogenesis and oncolysis.

### oHSV+trabectedin synergy extends to immunocompetent osteosarcoma models

We also sought to determine the effect of trabectedin in immunocompetent tumor models. Because none exist for Ewing sarcoma, we utilized immunocompetent osteosarcoma mouse models to determine the effects of combination treatment in a model with an adaptive immune system. Mouse cells are relatively non-permissive to HSV-1 infection, thus necessitating the use of both human xenograft and murine tumor models to understand the effect of oHSV+trabectedin treatment in human patients – where HSV-sensitive human cells and a full immune system are both present.

We again observed the reproducibility of oHSV+trabectedin synergy in two osteosarcoma models, K7M2 and F420, in immunocompetent mice (Figure 3A, B). Unlike the xenograft Ewing sarcoma models, neither osteosarcoma model responded to the monotherapies. Combination-treated animals possessed the lowest average tumor burden and highest response rates. In the case of K7M2, the combination treatment induced tumor regressions by day 28 (CR + PR = 8/8, 100%) and conferred a significant survival advantage (Figure S1, S2). Unfortunately, we found that the combination was hepatotoxic in F420-bearing animals due to the C57BL/6 background and trabectedin dose alterations reduced efficacy (Figure S5A-F). This prevented us from assessing any potential combinatorial advantage in this model. However, before mice reached the toxicity-induced weight loss endpoint, the combination-treated F420 animals exhibited decreased tumor burden and high response and disease stabilization rates (CR + PR = 4/6, 67%; DSR = 6/6, 100%). Thus, osteosarcoma immunocompetent models maintained the efficacy of combination therapy observed in Ewing sarcoma xenograft models.

**Figure 3.**
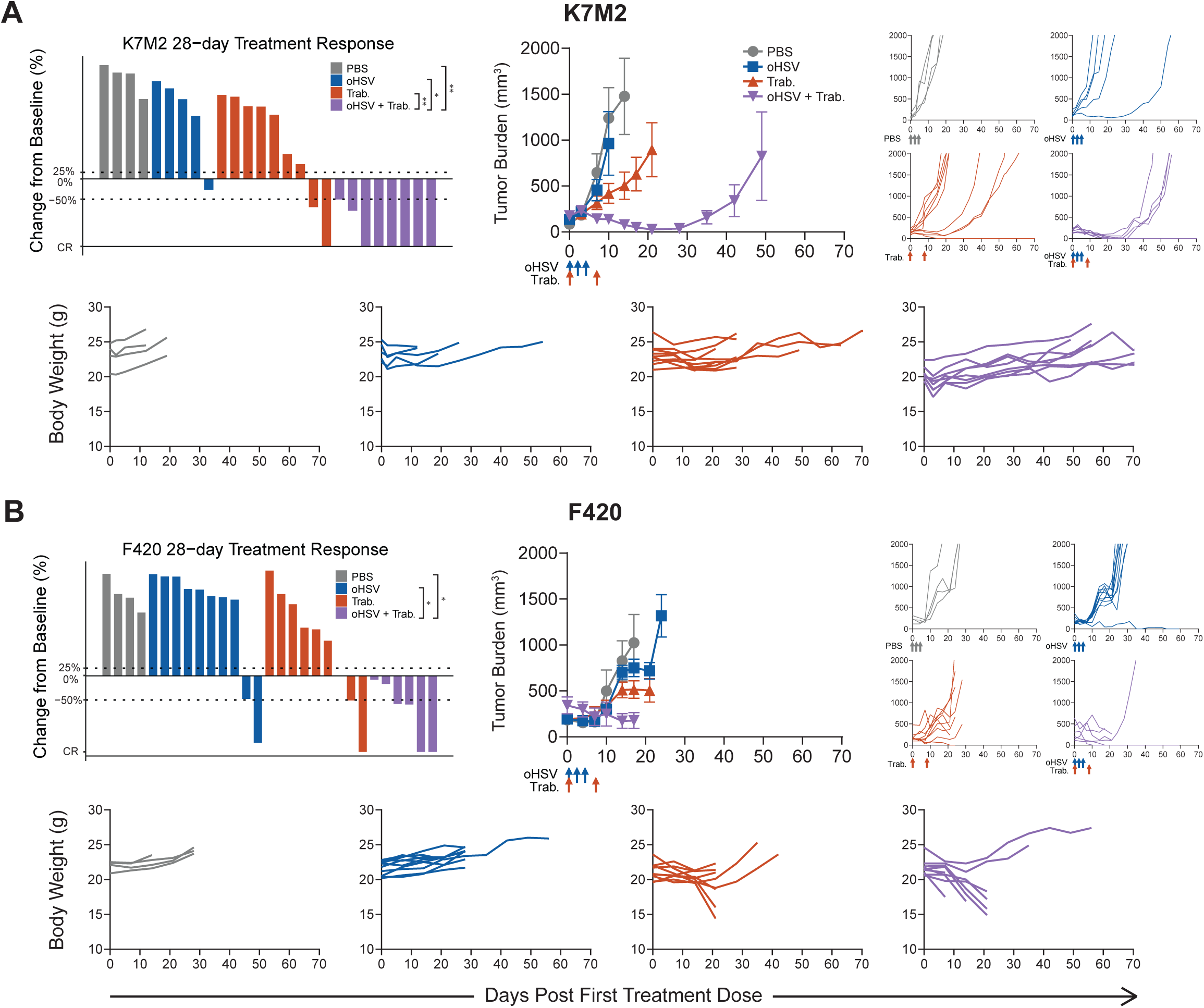
Trabectedin improves oHSV antitumor efficacy with reproducibility in immunocompetent osteosarcoma mouse models. The best response for each treated tumor through 28 days, the average tumor burden, spider plots, and individual body weights are shown for treated osteosarcoma models A) K7M2 and B) F420 in immunocompetent BALB/c and B6-albino mice, respectively. Body weight plots include mice that were excluded from the tumor burden analyses due to early non-tumor-related endpoints. PBS and oHSV (1.0 x 10^8^ pfu) were given intratumorally (i.Tu.) on Days 0, 2, 4. Trabectedin (0.15 mg/kg) was given intravenously (I.V.) on Days 0 & 7. Statistical analyses of the disease control rates (CR + PR + SD) were performed using a pairwise Fisher’s exact test with p values adjusted using the Benjamini-Hochberg procedure; *p < 0.05, **p < 0.01, ***p < 0.001. Summarized data with error bars depict mean ± SEM.

### Trabectedin modulates immunosuppressive cells and enhances cytotoxic effector stimulation

To determine the mechanism of combination synergy in immunocompetent osteosarcoma models, we again performed scRNAseq on treated tumors. Given the low infectivity of HSV-1 for mouse cells, we focused on other potential trabectedin-induced changes. Characterization of all cells in the microenvironment using mouse cell type references and cell marker gene expression coupled with unbiased recursive cluster assignment showed infiltration of NK cells, T cells, monocytes, and macrophages (Figure 4A, Figure S6A-D). Pathway analysis for each of the infiltrating cell types indicated an enrichment of the KEGG pathway “Natural killer cell mediated cytotoxicity” in infiltrating NK and T cells (Figure 4B). Using florescence-associated cell sorting (FACS) to complement these observations on a protein level, we noted that a significant increase in antitumor GP70^+^ CD8 T cell numbers occurred in combination-treated tumors, despite no significant combinatorial increase in overall CD8 or NK cell counts (p ≤ 0.05; Figure 4C, Figure S7A-B). We also observed a decrease in immunosuppressive cells, including M2 macrophages, T regulatory cells, and CTLA4^+^ T cells, following treatment (Figure 4C).

**Figure 4.**
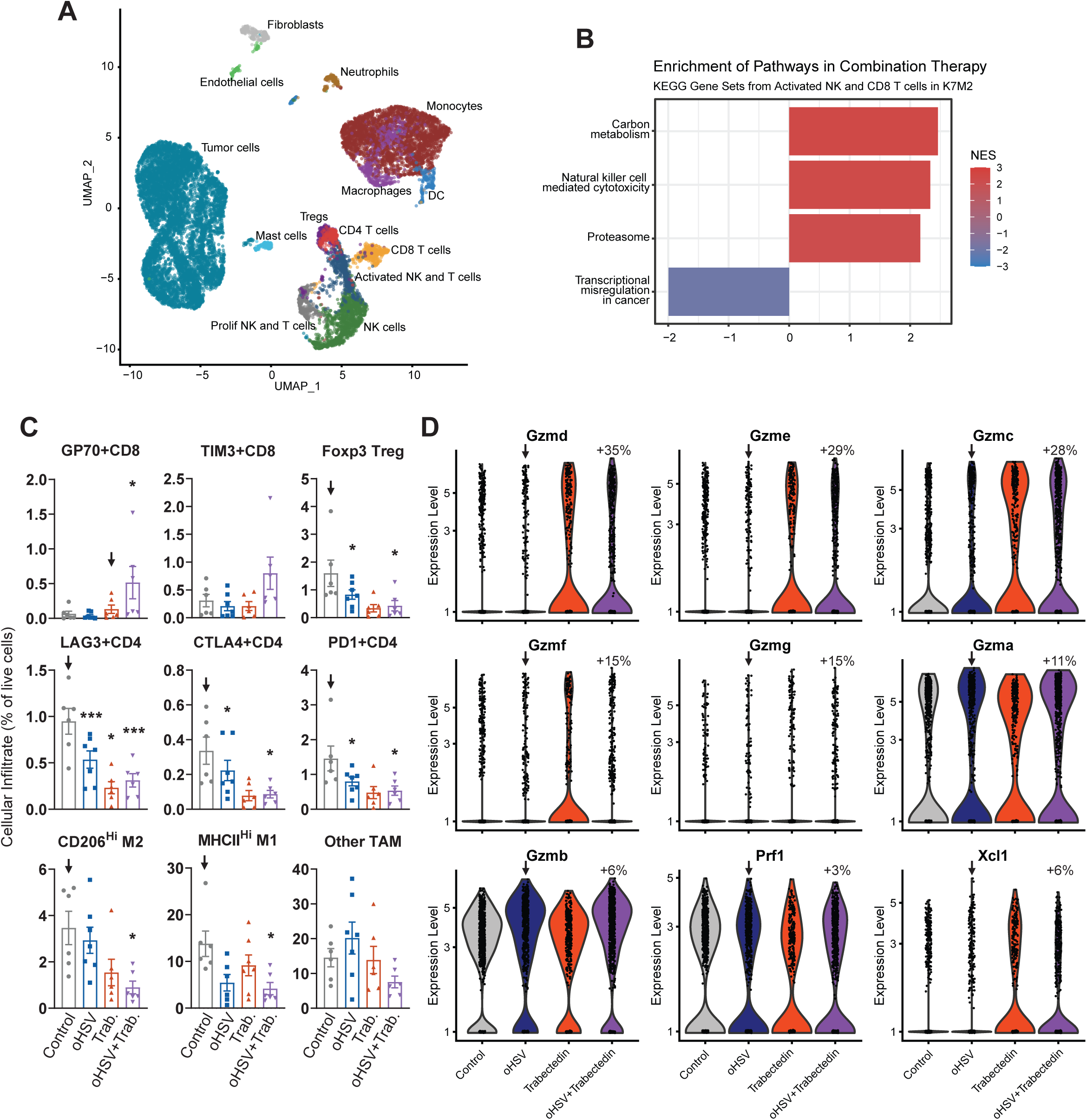
Trabectedin reduces immune suppression and enhances effector cell activation to synergize with oHSV. A) scRNAseq UMAP plot of the treated K7M2 tumor microenvironments. Datasets from all treatment groups were merged with no need for batch correction. B) Bar plot displaying the normalized enrichment score (NES) of the significant KEGG pathways from gene set enrichment analysis in NK cells from the tumor microenvironment. C) FACS of tumor infiltrating immune cells showed decreased immunosuppressive cells and increased activated T cells post-treatment (arrow indicates control for comparison, *p < 0.05, **p < 0.01, ***p < 0.001). D) Violin plots showing increased cytotoxic expression in all NK and T cells of combination-treated K7M2 (arrow indicates control for comparison, percent fold change = 100*2log2FC). PBS and oHSV (1.0 x 10^8^ pfu) were given i.Tu. on Days 0 & 2. Trabectedin (0.15 mg/kg) was given I.V. on Day 0. scRNAseq samples were collected on Day 3. Flow cytometry samples were collected on Day 7.

To investigate whether these patterns of enriched effector lymphocytes and decreased suppressive cells correlated with an activation of effector lymphocytes, we isolated NK and T cells from the scRNAseq datasets and probed for phenotypic changes between treatment groups. We found that trabectedin stimulated the expression of all granzyme (*Gzm*) genes, markers of cytotoxic activation (Figure 4D). Perforin (*Prf1*) expression was maintained across treatment groups. The combinatorial therapy also enhanced the potential activation of dendritic cells through increased expression of *Xcl1* by NK cells, a factor known to facilitate dendritic cell activation of T cells.^35–37^ Trabectedin promoted an increase in a cytotoxic lymphocyte phenotype that may induce immune-mediated tumor cell death.

### NK and CD8 T cells mediate combination efficacy via trabectedin-stimulated cytotoxicity in immunocompetent models

We conducted loss-of-function experiments to determine the reliance of combinatorial efficacy on NK or T cells individually. We first analyzed K7M2 osteosarcoma tumor burden and treatment responses in the immune-compromised (T and B cell-deficient) athymic nude mouse model. We found that the combinatorial efficacy was significantly diminished in this immunodeficient model – only one combination-treated tumor responded (CR + PR = 1/8, 13%). Similarly, we observed a higher average tumor burden (Figure 5A) than that observed in the immunocompetent K7M2 model (Figure 3A). However, the individual tumor growth curves revealed that the combination-treated tumors progressed slower than their monotherapy-treated counterparts. This maintained, albeit lessoned, synergy correlated with a significant survival advantage for the combination-treated nude mice (p = 0.0135; Figure S2). A study of F420 in nude mice resulted in similar efficacy outcomes (Figure S5F). These observations suggest a role for T cells in combination-induced regressions, however, the lack of T cells alone was insufficient to completely lose a combinatorial effect.

**Figure 5.**
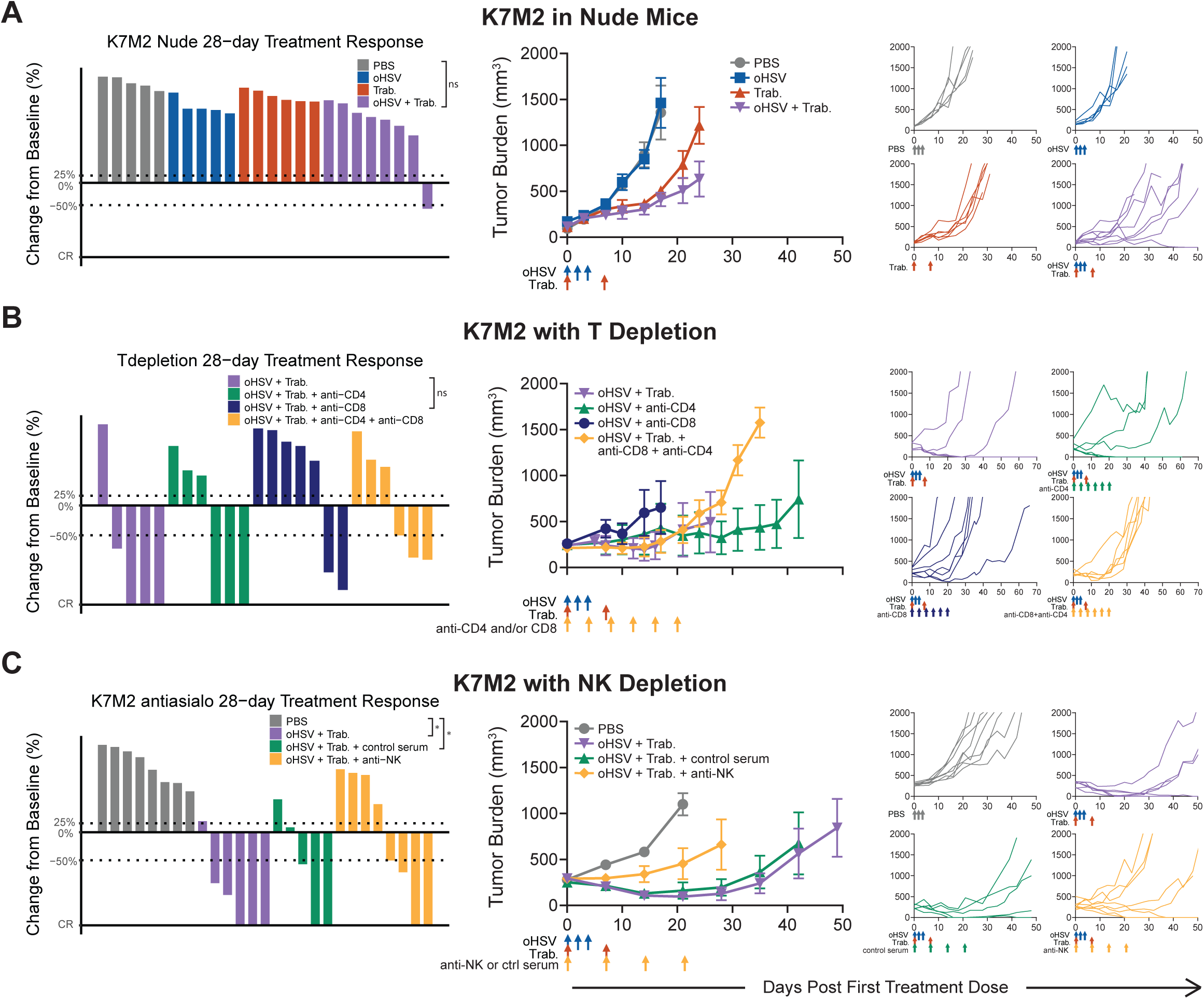
Combination efficacy in immunocompetent osteosarcoma models results from augmentation of antitumor NK and T cell responses. The best response for each treated tumor through 28 days, the average tumor burden, and spider plots tracking individual tumor volumes over the full study period are displayed for A) K7M2 tumor-bearing nude mice (lack T and B cells), B) K7M2-bearing BALB/c mice treated with anti-asialo NK cell depletion (given I.V. on Days 0, 7, 14, 21), C) K7M2-bearing BALB/c mice treated with antibody-mediated CD4 and CD8 T cell depletion (given I.V. on Days 0, 4, 8, 12, 16, 20, 24, 28). PBS and oHSV (1.0 x 10^8^ pfu) were given i.Tu. on Days 0, 2, & 4. Trabectedin (0.15 mg/kg) was given I.V. on Days 0 & 7. Statistical analyses of the disease control rates (CR + PR + SD) were performed using a pairwise Fisher’s exact test with p values adjusted using the Benjamini-Hochberg procedure; *p < 0.05, **p < 0.01, ***p < 0.001. Summarized data with error bars depict mean ± SEM.

We next performed antibody-mediated depletion of cytotoxic lymphocytes in treated mice with K7M2 osteosarcoma tumors. To further investigate the role of T cells in treatment efficacy, we depleted CD8 and CD4 T cells individually and concurrently (Figure 5B). Depletion of CD8 T cells conferred a loss of efficacy compared to combination-treated tumors with a complete immune system (CR + PR = 2/7, 29%). When we depleted CD4 T cells but left the CD8 subset intact, combinatorial therapy achieved several maintained complete responses (CR = 3/6, 50%). Thus, removal of CD4 T cells further increased the combinatorial efficacy. Indeed, when we analyzed the CD4 T cell subset of the scRNAseq dataset, a large proportion of the infiltrating CD4 T cells possessed a T regulatory cell phenotype (Figure 4A, Figure S6C, D). Finally, the depletion of both CD8 and CD4 T cells maintained initial responses through day 28, although tumors progressed quickly thereafter (PR = 3/6, 50%).

NK cell depletion caused treated animals to lose combinatorial efficacy, indicating that NK cell effector function is critical in the mechanism of oHSV and trabectedin synergy (Figure 5C). In NK-depleted animals, the combinatorial response rate decreased and the average tumor burden increased (CR + PR = 4/8, 50%). This response rate was similar to that of T cell-depleted mice, suggesting that both cell types contribute to oHSV+trabectedin synergy. The role of NK cells in combinatorial efficacy may indicate why mice lacking T cells maintained some combinatorial therapy effects. Thus, an oHSV-stimulated immune infiltration synergized with the trabectedin-mediated increase in cytotoxic NK and T cells to cause tumor regressions in immunocompetent osteosarcoma models.

## Discussion

Herein we report the combination of oHSV and trabectedin demonstrated generalizable synergy across human and mouse preclinical tumor models of pediatric bone sarcomas. Our data support a mechanism where oHSV and trabectedin synergize by enhanced oHSV spread in tumor cells and increased effector lymphocyte cytotoxicity. Trabectedin disrupts the expression of the intrinsic antiviral response within human tumor cells, leading to increased oHSV cellular pathogenesis. Additionally, trabectedin increases the intratumoral presence of antitumor (GP70^+^) CD8 T cells and stimulates the expression of granzyme genes in NK and T cells to achieve cytotoxic lymphocyte-dependent tumor regressions. Through these synergistic mechanisms, trabectedin amplifies the antitumor activities of oHSV against pediatric bone sarcomas.

oHSV targets tumor cells through two mechanisms – a tumor-specific infection that leads to oncolysis and immune recruitment and activation against tumor cells. Our studies demonstrate that trabectedin may augment both mechanisms, likely through its ability to bind DNA and disrupt gene expression. While our original rationale for the combination assumed an indirect effect of trabectedin on effector cells via a reduction in immunosuppressive monocytes and macrophages, our results suggest direct and indirect effects. Trabectedin reduced enrichment of the virus-induced intracellular antiviral response in human Ewing sarcoma cells. This decrease in antiviral expression coincided with an increase in oHSV transcript spread and intracellular abundance. We note that increased oHSV transcript spread does not exclude the possible transfer of oHSV transcripts by extracellular vesicles.^38,39^ However, we maintain that trabectedin increased oHSV transcript spread and intracellular abundance and that these effects were associated with tumor regressions.

In immunocompetent osteosarcoma, trabectedin increased cytotoxic gene expression in the presence and absence of oHSV – suggesting an effect on NK and T cells despite minimal oHSV-stimulated myeloid recruitment (Figure 4D, Figure S6B). While this observation does not exclude the possibility of myeloid targeting by trabectedin, the data indicate possible additional effects on NK and T cell cytotoxicity. Our results are consistent with other studies that reported increased T and NK cell activation and cytotoxicity following trabectedin treatment.^23,40,41^ Trabectedin is known to deplete and modulate the immunosuppression of myeloid cells and thereby indirectly lead to greater antitumor immune activity.^20,22^ Outside of an impact to monocytes and macrophages, trabectedin may indirectly enhance NK and T cell activity through signaling resulting from DNA damage and cellular stress in tumor cells.^40^ Alternatively, trabectedin may exert a direct effect on NK and T cells through an unexpected mechanism to stimulate cytotoxicity. Given the broad mechanisms of action by trabectedin and the various effects we observed herein, our data are consistent with the notion that a combination of the above mechanisms may activate cytotoxic lymphocytes.

We acknowledge that our preclinical models lack the ability to recapitulate a human patient tumor. We aimed to capture the direct antitumor and immune-mediated aspects of the patient response to the combinatorial therapy by employing human xenograft tumor models in immunodeficient mice to understand the effects on virus infection and oncolysis as well as immunocompetent mouse tumor models to elucidate the impacts on antitumor immunity. Given the lack of a mouse model of Ewing sarcoma, all Ewing sarcoma experiments necessitated implantation of human Ewing sarcoma cells into an immunodeficient mouse.^42^ While this enabled us to study the spread of oHSV in human Ewing sarcoma cells, we were limited in our ability to study a competent immune response. These models not only lack T and B cells, but NK-tumor interactions may be disrupted due to species-dependent differences in ligand-receptor interactions.^43^ In order to better clarify the role of the innate and adaptive immune response to our combinatorial therapy, we employed an immunocompetent mouse osteosarcoma model. This model was important because NK and T cell interactions may be essential to achieve immune-dependent efficacy. Evidence suggests that NK cells stimulate dendritic cell recruitment and their downstream activation of T cells.^37^ The triad of NK cell, dendritic cell, and T cell interactions occurring in an immunocompetent model may be critical for immune-mediated combinatorial efficacy. By using the osteosarcoma syngeneic model, which has both innate and adaptive immunity, we captured the immune-related synergistic mechanisms of oHSV+trabectedin.

It is possible the use of different histologic models may, in part, determine the effects of the combination therapy. Osteosarcoma may be predisposed to the DNA-damaging effects of trabectedin which induce stress signals that lead to NK and T cell activation.^41^ Ewing sarcomas possess unique sensitivity to trabectedin, since trabectedin targets fusion oncogenes and displays efficacy as a monotherapy in tumors with fusion oncogene dependencies.^26,28^ Future work could address these discrepancies through a validation of the combinatorial synergistic mechanisms in additional human and mouse tumor lines and humanized mouse models. Recent strides have been made in developing an immunocompetent Ewing sarcoma model.^44,45^ Future investigations would benefit from utilizing such approaches to cohesively capture the tumor intrinsic and immune-dependent mechanisms of oHSV+trabectedin synergy.

Our findings come at an encouraging time in the field, where recent preclinical and clinical results highlight the therapeutic potential for trabectedin. Preclinical and clinical data demonstrated Ewing sarcoma vulnerability to fusion oncogene (EWS-FLI or other FET/ETS fusion) targeting by trabectedin.^26,28–31,46^ These studies and others underscored the clinical relevance of trabectedin against Ewing sarcoma, despite initial challenges to achieve clinical outcomes at lower serum concentrations.^28,47^ Indeed, a recent multicenter phase I study of trabectedin as a one-hour infusion in combination with irinotecan in recurrent Ewing sarcoma displayed clinical benefit and safety in pediatric patients, providing rationale for an ongoing phase II study.^33^ While trabectedin can induce grade 3 and 4 hepatotoxicity, which we also observed in the C57BL/6 mouse background, the drug is FDA-approved and considered safe in patients, including pediatric patients^19,31,32^. The FDA approval of T-VEC and the evidence of oHSV clinical safety in pediatric patients corroborate the clinical application of oHSV+trabectedin combination therapy.^2,17,18^ The combination of trabectedin and oHSV capitalizes on the direct effects of trabectedin on Ewing sarcoma and strengthens tumor cell stress and immune involvement. A recent phase II clinical trial displaying synergy between T-VEC, trabectedin, and nivolumab in advanced sarcomas supports the synergistic results that we observed and provides further clinical rationale for combining oHSV and trabectedin to treat pediatric patients with bone sarcoma.^48^

Our work demonstrates the synergy of oHSV and trabectedin in preclinical pediatric bone sarcoma models and clarifies the tumor intrinsic and immune-mediated mechanisms whereby trabectedin augments viroimmunotherapy. Our future work will continue to focus on understanding these synergistic mechanisms by examining the combinatorial effects in additional models of pediatric bone sarcoma and moving towards clinical application. The combination of oHSV and trabectedin could target tumors in pediatric patients and provide further rationale for synergistic combinations, offering a promising opportunity to improve outcomes for children, adolescents, and young adults with bone sarcoma.

## Materials and Methods

Our objective of this study was to determine if the enhanced antitumor efficacy observed when combining oHSV with trabectedin to treat human EWS xenograft tumor models is generalizable over a panel of PDXs and immunocompetent murine tumor models using oHSV variants. We aimed to elucidate the mechanisms of synergy in both immunodeficient and immunocompetent tumor models.

### Cell Lines, Patient-derived Xenografts (PDXs) and Viruses

#### Cell lines

The A673 human Ewing sarcoma cell line (Cat# CCL-81), Vero green monkey kidney cell line (Cat# CRL-1598), and K7M2 mouse osteosarcoma cell line (Cat# CRL-2836) were purchased from the American Type Culture Collection (ATCC) (Manassas, VA). K7M2 was derived from a spontaneous metastatic lesion in a female BALB/c background.^49^ F420 was originally derived from a female C57BL/6 mouse transgenic for a mutant p53 under the control of a bone-selective promoter (kind gift from Jason Yustein).^50^ The identities of cell lines were confirmed by short tandem repeat (STR) genotyping. Mycoplasma testing was performed yearly via PCR. A673, K7M2, F420, and Vero cell lines were cultured in Dulbecco’s Modified Eagle Medium (DMEM) supplemented with 10% fetal bovine serum (FBS), 200 IU/mL penicillin and 200 µg/mL streptomycin (Gibco brand - Thermo Fisher Scientific, Waltham, MA). New stocks of cell lines were thawed every five to eight passages *in vitro* (approximately 1 month).

#### PDXs

PDXs CHLA258 and EW5 (both Ewing sarcoma) and PDX-0027 (rhabdomyosarcoma) were obtained as frozen vials of tissue fragments (∼2 mm x 2 mm) from the Tumor Core at Nationwide Children’s Hospital (Columbus, OH). The thawing and surgical implantation of PDX tissues are described in the Animal Studies section.

### Viruses

rRp450 (strain KOS), attenuated by insertion of the rat cytochrome p450 CYP2B1 gene into the viral ribonucleotide reductase ICP6 gene, was provided by E. Antonio Chiocca (Brigham and Women’s Hospital, Boston, MA).^51^ HSV1716 (strain 17), attenuated by a deletion in RL1 that encodes ICP34.5, was provided by Virttu Biologics (Glasgow, UK; later acquired by Sorrento Therapeutics, San Diego, CA) and maintained and propagated as previously described.^52^

### Compounds and Reagents

Trabectedin was procured from two sources: (1) a generous gift from Janssen Pharmaceutical Companies of Johnson and Johnson (Titusville, NJ) via a materials transfer agreement and (2) purchased from MedChem Express (Monmouth Junction, NJ). For *in vivo* studies, trabectedin provided by Janssen Pharmaceutics was formulated at 0.0375 mg/mL in sterile purified water. Trabectedin purchased from MedChem Express was dissolved in DMSO at 4 mg/mL and stored at -20°C. On the day of injection, trabectedin was thawed and formulated in 0.1 g of Hydroxy B-Cyclodextrin (Sigma-Aldrich, St. Louis, MO, MKBL6868V) and sterile PBS at 0.0533 mg/mL. For all studies, trabectedin was injected in a final volume of 100 µL per mouse at 0.15 mg/kg. Dexamethasone was purchased at 4 mg/mL from Mylan Pharmaceuticals (Morgantown, WV) and diluted in PBS. Lyophilized anti-asialo GM1 antibody (anti-NK) in normal rabbit serum was obtained from FujiFilm Wako Pure Chemical (Richmond, VA) and dissolved in 1 mL sterile distilled water. Frozen sterile normal rabbit serum was acquired from Thermo Fisher Scientific.

### Animal Studies

All animal studies were approved by the Nationwide Children’s Hospital Institutional Animal Care and Use Committee under protocols AR12-00074 and AR19-00116.

#### Mice

Athymic nude (Hsd:Athymic Nude-*Foxn1^nu^)*, BALB/c (BALB/cAnNHsd), C57BL/6, and B6-albino (B6(Cg)-Tyrc-2J/J) female mice were purchased from Envigo (Indianapolis, IN). CB17 (Scid C.B-*Igh-1^b^*/IcrTac-*Prkdc^scid^*) female mice were purchased from Taconic (Germantown, NY). NSG-SGM3 (NOD.Cg-*Prkdc^scid^ Il2rg^tm1Wjl^* Tg(CMV-IL3,CSF2,KITLG)1Eav/MloySzJ) female mice were purchased from Jackson Labs (Bar Harbor, ME). All mice arrived at 4-5 weeks of age and were enrolled in animal studies at 6-8 weeks of age. Humane endpoints included a tumor volume of 2000 mm^3^, a tumor length of 2 cm, 20% weight loss, treatment-related ulceration (> 1 cm) or blistering/necrosis, or a body condition score of less than 2. Mice that reached a non-tumor-related endpoint prior to sufficient data collection (2.5 weeks) were excluded from the study.

#### PDX Tumor Models

After thawing frozen vials at 37°C, the PDX tumor cell line A673 was passaged at least twice in culture to reach exponential phase and then implanted by simple subcutaneous injection (5.0x10^6^ A673 tumor cells) in the right flank of female CB17S or athymic nude mice. PDX tissue fragments were thawed and immediately rinsed with warm DMEM (37°C) and surgically implanted subcutaneously (2 x 2 mm^3^ tumor fragment) in the right flank CB17S (PDX-0027) or athymic nude mice (CHLA258, EW5). Tumors derived from the PDX tissue fragments were allowed to grow to a volume of ∼1800 mm^3^ and then donor mice were humanely euthanized. After the tumors derived from PDX tissues were harvested and diced into 2 x 2 mm^3^ tumor fragments, these fresh tumor fragments were surgically implanted subcutaneously in the right flank of CB17S (PDX-0027) or athymic nude (CHLA258, EW5) female mice. Both PDX cell line-derived and PDX tissue-derived tumors were allowed to reach ∼200 mm^3^ prior to enrollment into the efficacy studies. Mice-bearing PDX tumors were then treated with trabectedin intravenously (I.V.) by tail vein injection (Days 0 and 7). rRp450 or HSV1716 was diluted in PBS to 1.0 x 10^7^ plaque-forming units (pfu)/mL per mouse (each mouse receiving 100 µL) and administered intratumorally (i.Tu.) on Days 0 and 2. PBS (100 µL per mouse) was administered i.Tu. as the control for rRp450 or HSV1716 (Days 0 and 2).

#### Murine Tumor Models in Immunocompetent Mice

After thawing, F420 or K7M2 tumor cell lines were passaged at least twice in culture to reach exponential phase and prior to implantation. F420 (derived from female C57BL/6 mice) was implanted in either C57BL/6 or B6-albino (C57BL/6 background) female mice and K7M2 (derived from female BALB/c) in female BALB/c or athymic nude (BALB/c background) mice. F420 was implanted in the right flank by simple subcutaneous injection (5.0 x 10^6^ tumor cells). K7M2 was initially implanted in the right flank of female athymic nude mice by simple subcutaneous injection (2.5 x 10^6^ tumor cells). To enhance K7M2 growth, K7M2 tumors were allowed to grow to a volume of ∼1800 mm^3^ and then donor mice were humanely euthanized. After the large K7M2 tumors were harvested and diced into 2 x 2 mm^3^ tumor fragments, these tumor fragments were surgically implanted subcutaneously in the right flank of female BALB/c mice. These mice then entered studies or became K7M2 donor mice for future studies. Efforts were made to keep tumor passages in the immunocompetent BALB/c mice low and to use tumor passages within several months on the same study. Then, mice bearing F420 or K7M2 flank tumors (∼150-300 mm^3^) were treated with trabectedin I.V. by tail vein injection or retro-orbital injection (Days 0 and 7). HSV1716 was diluted in PBS to 1.0 x 10^8^ pfu/mL per mouse (each mouse receiving 100 µL) and administered i.Tu. (Days 0, 2, and 4). PBS (100 µL per mouse) was administered i.Tu. as the control for HSV1716 (Days 0, 2, and 4).

#### Murine Tumor Models in Immunodeficient Mice

K7M2 tumor cell lines were passaged at least twice in culture to reach exponential phase and then implanted by simple subcutaneous injection (2.5 x 10^6^ tumor cells) in the right flank of female athymic nude mice. Athymic nude mice bearing K7M2 flank tumors (150-200 mm^3^) were then treated with trabectedin I.V. by retro-orbital injection (Days 0 and 7). HSV1716 was diluted in PBS to 1.0 x 10^8^ pfu/mL per mouse (each mouse receiving 100 µL) and administered i.Tu. (Days 0, 2, and 4). PBS (100 µL per mouse) was administered i.Tu. as the control for HSV1716 (Days 0, 2, and 4).

#### oHSV Replication Studies

A673 xenograft tumors from mice treated with HSV1716 or HSV1716+Trabectedin regimens previously described were harvested at 72 hours and 144 hours after infection. Tumors were homogenized and subjected to three cycles of freezing and thawing before a 5–8 log serial dilution in 10% FBS DMEM and an inoculation of a 90% confluent 12-well plate of Vero cells. The virus was applied for 2 hours before adding overlay media (1% carboxymethylcellulose [CMC], 1xMEM, 10%FBS) and incubating in a 37°C incubator for 2 days. Plates were stained in crystal violet for 20 minutes, washed in tap water, and dried overnight. Viral plaques were counted and multiplied by serial dilution to determine viral titer (pfu).

#### Single-cell RNA sequencing

A673 (n = 8) and K7M2 (n = 4, bi-flank) tumors were implanted into the flanks of athymic nude and BALB/c mice, respectively, according to the methods described above for each model. Mice bearing tumors ∼300-500 mm^3^ were selected for the scRNAseq experiments. Mice were given trabectedin I.V. by tail vein injection (Day 0). HSV1716 was diluted in PBS to 1.0 x 10^7^ pfu/mL (A673) or 1.0 x 10^8^ pfu/mL (K7M2) per mouse (each mouse received 100 µL) and administered i.Tu. on Days 0 and 2. PBS (100 µL per mouse) was administered i.Tu. on Days 0 and 2.

To create single cell suspensions, tumors were harvested from mice on Day 3 and separately dissociated on a GentleMacs Octo Dissociator with Heaters (Miltenyi Biotec, Bergisch Gladbach, Germany, 130-096-427) using the human tumor dissociation kit (Miltenyi Biotec, 130-095-929). Dead cells were removed using the dead cell removal kit (Miltenyi Biotec, 130-090-101) per the manufacturer protocol. Viable single cell suspensions were resuspended in 0.04% BSA in PBS (Sigma-Aldrich, A1595-50ML and Gibco, 21-031-CV) and loaded onto the Chromium Next GEM Chip G Single Cell Kit (10x Genomics, PN-1000120) at the appropriate volume required to target 5000-8000 cells per sample. Final libraries were loaded onto an Illumina NovaSeq 6000 sequencer to generate a minimum of 65 million reads per sample.

#### CD4^+^ and CD8^+^ Cell Depletion with Monoclonal Antibodies

To deplete CD4^+^ and/or CD8^+^ lymphocytes, K7M2 bearing female BALB/c mice were established as described above in the efficacy studies and then administered IP with either 250 μg of anti-CD4 antibody (GK1.5), 250 μg of anti-CD8 antibody (YTS169.4) or both at 250 μg of anti-CD4 and CD8 antibodies (Days 0, 4, 8 and 12). Mice were injected i.Tu. with either 100 μL of PBS (control) or 1.0 × 10^8^ pfu HSV1716 in 100 μL of PBS on Days 0, 2 and 4. Trabectedin was administered I.V. by retro-orbital injection on Days 0 and 7. The oHSV+trabectedin cohort was derived from a later K7M2 tumor passage than the T cell-depleted cohorts, but shared the same initial athymic nude donor cohort.

#### NK Cell Depletion with Anti-Asialo GM1

To deplete NK cells, K7M2-bearing female BALB/c mice were established as described above in the efficacy studies and then administered IP with 50 μL anti-asialo GM1 antibody (FujiFilm Wako Pure Chemical, Richmond, VA) or control rabbit serum (Thermo Fisher Scientific) on Days 0, 7, 14 and 21. Mice were injected i.Tu. with either 100 μL of PBS (control) or 1.0 × 10^8^ pfu HSV1716 in 100 μL of PBS on Days 0, 2 and 4. Trabectedin was administered I.V. by retro-orbital injection on Days 0 and 7.

#### Fluorescence-Activated Cell Sorting (FACS) Analysis

K7M2 tumors (∼400 mm^3^) were established in BALB/c mice as described in the efficacy studies. Mice were then treated with trabectedin I.V. by retro-orbital injection (Day 0). HSV1716 was diluted in PBS to 1.0 x 10^8^ pfu/mL per mouse and administered i.Tu., as previously described (Days 0, 2, and 4). PBS was administered i.Tu. as the control for HSV1716, as previously described (Days 0, 2, and 4). Animals were then euthanized on Day 7 and whole tumors collected and transferred into 6-well dishes with 2 mL 1xPBS. Samples were minced by mechanical chopping and incubated in 1xPBS containing 150 mg/mL DNAse I (Roche Diagnostics, Indianapolis, IN) and 25 mg/mL liberase (Roche Diagnostics) at 37°C for one hour. Tumor samples were then filtered through 70 µM cell strainers (Thermo Fisher Scientific). Single-cell suspensions were spun down and tumor cells were lysed with ACK RBC lysis buffer (Lonza Group, Basel, Switzerland) at room temperature for 5 minutes. Tumor cells were washed with 1xPBS and blocked with 5% mouse Fc blocking reagent (Biolegend, San Diego, CA) in FACS buffer (1% FBS and 1 mM EDTA in PBS) on ice for 30 minutes.

Tumor samples were labeled on ice for 30 minutes with one of the following antibody staining panels. All of the following flow cytometry antibodies were purchased from BioLegend (San Diego, California) except for anti-CD25 (BD Biosciences, San Jose, CA), anti-Ly6C (BD Biosciences), and anti-Foxp3 (eBiosciences, San Diego, CA). For NK and lymphocytes: CD4-fluorescein isothiocyanate (FITC) (GK1.5), CD49b-phycoerythrin (PE) (DX5), CD8a-PE-Cy7 (53-6.7), B220-PerCP/Cy5.5 (RA3-6B2), CD44-allophycocyanin (APC) (IM7) and CD3e-Violet 421 (145-2C11). For T cell exhaustion markers: CD4-FITC, CD8a-PE-Cy7, B220-PerCP/Cy5.5 and CD3e-Violet 421 together with LAG3-PE (C9B7W) and Tim3-APC (RMT3-23) or CTLA4-PE (UC10-4B9) and PD-1-APC (29F.1A12). For myeloid cells: CD11b-Violet 421 (M1/70), F4/80-PE-Cy7 (BM8), Ly6C-APC (AL-21), Ly-6G-APC-Cy7 (1A8); For GP70^+^ CD8 T cells: GP70-TETRAMER-APC, mouse MuLV gp70 env 423 SPSYVYHQF, obtained from the NIH tetramer core facility (Human B2M H-2Ld). Labeled cells were washed with FACS buffer and fixed in 1% paraformaldehyde (Thermo Scientific Chemicals brand – Thermo Fisher Scientific) on ice for 5 minutes. CD25^+^Foxp3^+^ T regulatory cells were identified using intra-cellular staining. Mononuclear cells were enriched using Percoll (GE Healthcare, Little Chalfont, United Kingdom) density gradient centrifugation. Tumor single-cell suspensions were spun down. Cell pellets were suspended in 5 mL of 44% Percoll and gradually loaded on top of 3 mL of 67% Percoll in a 15 mL tube. Sample tubes were spun at 500 x g without brake at room temperature for 20 minutes. The interface cells between 44% and 67% Percoll were collected. Mononuclear cells were washed with FACS buffer and blocked with 5% mouse Fc blocking reagent in FACS buffer. Cells were stained with CD4-APC, CD8-PE-Cy7, CD25-PE, CD49b-PerCP, B220-APC-CY7, and CD3e-Violet 421 on ice for 30 minutes followed by Foxp3-FITC (FJK-16s) intra-cellular staining using cell fixation and cell permeabilization kit (Invitrogen). Stained cells were fixed in 1% paraformaldehyde. Data were collected on a BD FACS LSR II (BD Biosciences) and results were analyzed using the FlowJo software, version 10.0.3 (Tree Star Inc.).

#### Animal Studies Analysis and Plot Generation

Best overall response rates for each experiment were calculated within the treatment response period of 28 days using R. The following definitions were used: complete response (CR) – any tumor volume ≤ 0.10mm^3^ for at least one time point, partial response (PR) – relative tumor volume ≤ 0.5 for at least one time point, stable disease (SD) – does not meet the above definitions and has a relative tumor burden ≤ 1.25 at 28 days, progressive disease (PD) – does not meet the above definitions (relative tumor burden > 1.25 by 28 days). Relative tumor volume was defined as tumor volume divided by the initial tumor volume. The ggplot package was used to depict the relative tumor burden of the best overall response for each treated-tumor. GraphPad Prism v7.0 was used to plot the average tumor burdens (mm^3^), individual tumor burdens (mm^3^), and individual weights (g) for each treatment group. Once a mouse reached endpoint, plotting of the curve for that treatment group (for average tumor burden) or individual tumor burden ceased. GraphPad Prism v7.0 was also used to plot survival curves, cellular infiltrates, cytokine levels, toxicology factors of interest, and histological scoring.

### Single-cell RNA sequencing Analysis and Tools

#### Ewing sarcoma samples

Initial raw data processing, reference alignment, and feature-barcode matrix construction was performed via Cell Ranger (version 3.0.2, 10x Genomics) with reads mapped to the mm10 mouse genome (GCF_000001635.20), hg19 human genome (GCF_000001405.13), HSV-1 strain 17 genome (GCF_000859985.2) references. The count outputs were loaded into Seurat R toolkit V4 for further processing and analysis.^53^ The rrrSingleCellUtils package was used for early processing, quality control, and UMAP plots (*github.com/kidscancerlab/rrrSingleCellUtils*). Seurat quality control cutoffs were optimized for each sample, but cells with less than 800 RNA counts or greater than 40% mitochondrial reads were always omitted. Human cells (tumor cells) were isolated. All datasets were annotated, merged, and subsequently normalized (Seurat). Batch correction was not necessary. Principle component analysis, UMAP dimensional reduction, and unsupervised clustering (“FindNeighbors”, “FindClusters”, dims=1:18) were performed using Seurat. Differentially expressed genes and fold change values were identified using “FindMarkers” and “FoldChange” (Seurat, wilcox). HSV transcripts and the cells containing them were identified and visualized using “DimPlot”, and “VlnPlot” (Seurat, rrrSingleCellUtils). Module scores were calculated using the “AddModuleScore” function (Seurat) using HSV time point-dependent gene lists. Module scores were visualized with “FeaturePlot” and “VlnPlot” (Seurat, rrrSingleCellUtils). The clusterProfiler R package was used to perform gene set enrichment analysis using KEGG gene sets and the enrichplot R package provided the plotting via “gseaplot” for the “Herpes simplex virus 1 infection” KEGG gene set.^54,55^ The pathview package was used to depict the pathway of the “Herpes simplex virus 1 infection” KEGG gene set and the upregulated or downregulated genes within the pathway. The pathway results were summarized and depicted herein using Biorender.com. The ggplot2 package generically assisted with plot generation in R.

#### Osteosarcoma samples

Initial raw data processing, reference alignment, and feature-barcode matrix construction was performed using Cell Ranger (version 7.0.0, 10x Genomics) with reads mapped to the mm10 mouse genome (GCF_000001635.20), hg19 human genome (GCF_000001405.13), HSV-1 strain 17 genome (GCF_000859985.2) references. As above, the count outputs were loaded into Seurat R toolkit V4 for further processing and analysis and the rrrSingleCellUtils package was used for early processing, quality control, and UMAP plots. Seurat quality control cutoffs were optimized for each sample (individual cutoffs available with code on GitHub), but cells with less than 800 RNA counts or greater than 20% mitochondrial reads were omitted. All datasets were annotated, merged, and subsequently normalized (Seurat). Batch correction was not necessary. Principle component analysis, UMAP dimensional reduction, and unsupervised clustering (“FindNeighbors”, “FindClusters”, dims=1:17) were performed using Seurat. Unsupervised recursive sub-clustering was performed. Differentially expressed genes and fold change (log2FC) values were identified using the wilcox method for “FindMarkers” and “FoldChange” (Seurat). Percent increases in expression between groups were calculated from log2FC values using x%=(100*2^log2FC^-1). Cell type assignment was informed by SingleR assignment via the celldex references “MouseRNAseqData” (bulk RNAseq mouse reference) and “ImmGenData” (mouse immune reference).^56–58^ The reference annotation, recursive sub-cluster assignments, and most differentially expressed genes for each cluster (“FindMarkers”) were used to assign cell types.

The clusterProfiler R package was used to perform gene set enrichment analysis using KEGG gene sets and the enrichplot R package was used to depict the top significantly enriched results. The ggplot2 package assisted with plot generation.

### Statistical Analysis

Statistical tests were run using R or GraphPad Prism v7.0. Disease stabilization rates between groups underwent a pairwise to determine significant differences between treatment groups. Summarized data with error bars depict mean ± SEM. Survival studies were analyzed using the Mantel-Cox log-rank test. Otherwise, the comparison of samples with normal distribution was analyzed using Student’s t-test and the difference among group means was determined with ANOVA. Significant differences in sample groups were determined by p < 0.05.

## Data Availability Statement

The code utilized for the scRNAseq and response rate analyses herein is available on https://github.com/kidcancerlab/oHSV-trabectedin. scRNAseq data storage on GEO and

SRA is in progress.

## Acknowledgments

This work was generously supported by funding provided by the NIH/NCI, including the National Cancer Institute Cancer Moonshot Award U54CA232561, F31CA278353, T32CA269052, as well as CancerFree KIDS Foundation, Pelotonia Institute of Immuno-Oncology, Nationwide Children’s Director’s Strategic Development Fund, and CTSA Grant UL1TR002733. The content is solely the responsibility of the authors and does not necessarily represent the official views of our funding sources. We thank the Nationwide Children’s Hospital Tumor Core, Morphology Core, and the Steve and Cindy Rasmussen Institute for Genomic Medicine for their contributions to this work. BioRender.com was used to create the schemas within this manuscript.

## Author contributions

T.P.C., M.A.C., E.M.R., and R.D.R. conceived, designed, and directed the research and methodology. T.P.C. and R.D.R were co-advisors of E.M.R. on this project. D.A.L., E.M.R., P.J., E.S., and W.E.C. contributed to the conceptual development and evolution of the research.

M.A.C., A.M.G., and E.M.R. performed animal studies. C.Y.C. performed flow cytometry analyses. L.B. provided reagents for the depletion studies. P.Y.W. determined scRNAseq collection time point. M.C. and A.C.G. prepared next-generation sequencing libraries. E.M.R., M.V.C., and M.G. performed computational analyses. E.M.R., M.V.C., and R.D.R. reviewed computational analyses. L.E.B. stained and read immunohistochemistry slides. E.M.R. interpreted the consolidated data and wrote the manuscript. E.M.R., T.P.C., R.D.R., M.A.C., A.M.G., C.Y.C., M.V.C., M.C., A.C.G., M.G., P.Y.W., L.B., E.S., P.J., L.E.B., D.A.L., E.R.M., and W.E.C. edited the manuscript. E.M.R. revised the manuscript. All authors read and approved the final manuscript.

## Tables

None for the main manuscript body.

---

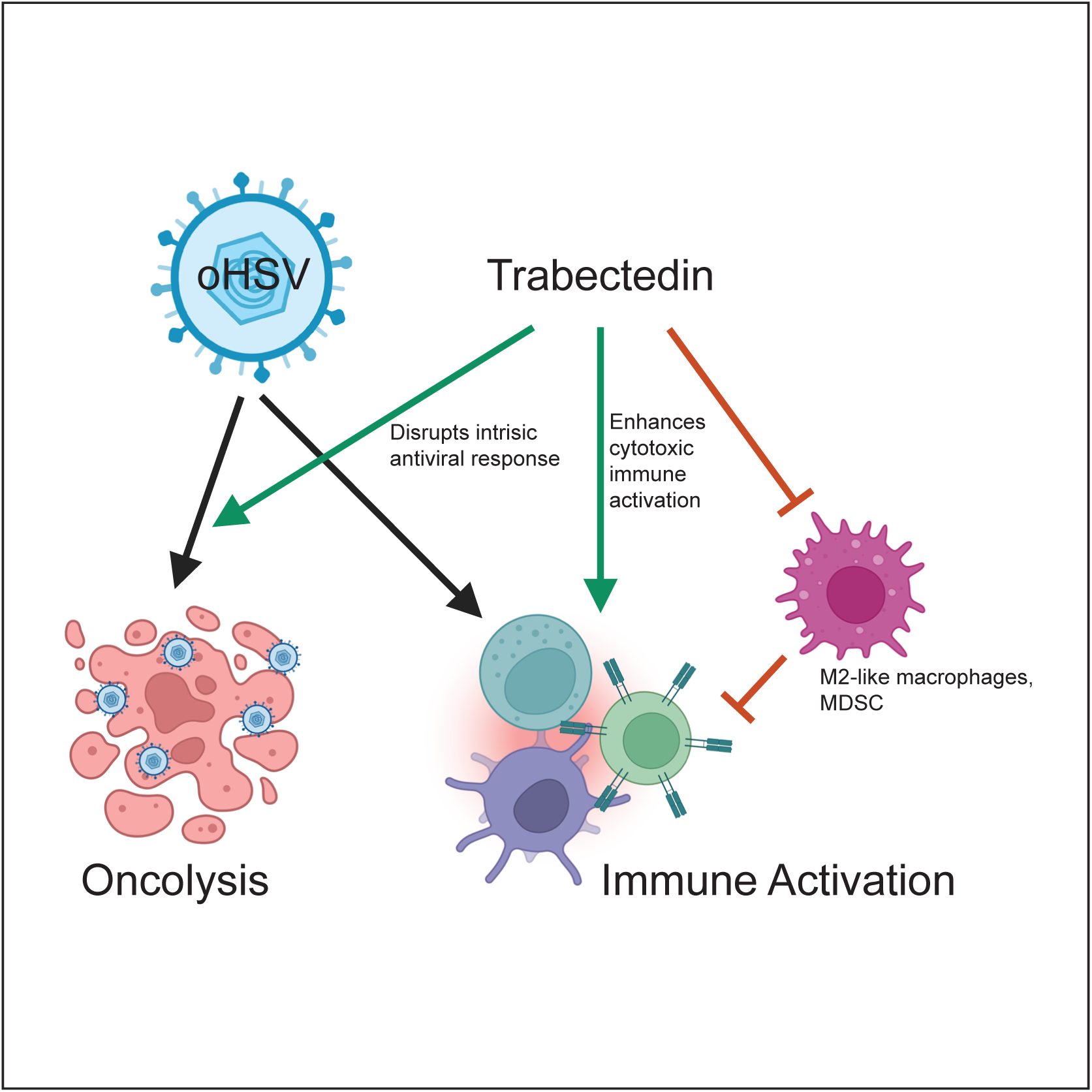

## References

1. Andtbacka, R.H., Kaufman, H.L., Collichio, F., Amatruda, T., Senzer, N., Chesney, J., Delman, K.A., Spitler, L.E., Puzanov, I., Agarwala, S.S., Milhem, M., et al. (2015). Talimogene Laherparepvec Improves Durable Response Rate in Patients With Advanced Melanoma. J Clin Oncol 33, 2780–2788. 10.1200/jco.2014.58.3377.

2. Haitz, K., Khosravi, H., Lin, J.Y., Menge, T., and Nambudiri, V.E. (2020). Review of talimogene laherparepvec: A first-in-class oncolytic viral treatment of advanced melanoma. J Am Acad Dermatol 83, 189–196. 10.1016/j.jaad.2020.01.039.

3. Shalhout, S.Z., Miller, D.M., Emerick, K.S., and Kaufman, H.L. (2023). Therapy with oncolytic viruses: progress and challenges. Nature Reviews Clinical Oncology 20, 160–177. 10.1038/s41571-022-00719-w.

4. Lin, D., Shen, Y., and Liang, T. (2023). Oncolytic virotherapy: basic principles, recent advances and future directions. Signal Transduction and Targeted Therapy 8, 156. 10.1038/s41392-023-01407-6.

5. Daw, N.C., Chou, A.J., Jaffe, N., Rao, B.N., Billups, C.A., Rodriguez-Galindo, C., Meyers, P.A., and Huh, W.W. (2015). Recurrent osteosarcoma with a single pulmonary metastasis: a multi-institutional review. Br J Cancer 112, 278–282. 10.1038/bjc.2014.585.

6. Van Mater, D., and Wagner, L. (2019). Management of recurrent Ewing sarcoma: challenges and approaches. Onco Targets Ther 12, 2279–2288. 10.2147/OTT.S170585.

7. Hamilton, S.N., Carlson, R., Hasan, H., Rassekh, S.R., and Goddard, K. (2017). Long-term Outcomes and Complications in Pediatric Ewing Sarcoma. Am J Clin Oncol 40, 423–428. 10.1097/COC.0000000000000176.

8. Dorfman, H.D., and Czerniak, B. (1995). Bone cancers. Cancer 75, 203–210. 10.1002/1097-0142(19950101)75:1+<203::aid-cncr2820751308>3.0.co;2-v.

9. Zarghooni, K., Bratke, G., Landgraf, P., Simon, T., Maintz, D., and Eysel, P. (2023). The Diagnosis and Treatment of Osteosarcoma and Ewing’s Sarcoma in Children and Adolescents. Dtsch Arztebl Int 120, 405–412. 10.3238/arztebl.m2023.0079.

10. Bernstein, M., Kovar, H., Paulussen, M., Randall, R.L., Schuck, A., Teot, L.A., and Juergens, H. (2006). Ewing’s sarcoma family of tumors: current management. Oncologist 11, 503–519. 10.1634/theoncologist.11-5-503.

11. Durfee, R.A., Mohammed, M., and Luu, H.H. (2016). Review of Osteosarcoma and Current Management. Rheumatol Ther 3, 221–243. 10.1007/s40744-016-0046-y.

12. Rajan, S., Franz, E.M., McAloney, C.A., Vetter, T.A., Cam, M., Gross, A.C., Taslim, C., Wang, M., Cannon, M.V., Oles, A., and Roberts, R.D. (2023). Osteosarcoma tumors maintain intra-tumoral transcriptional heterogeneity during bone and lung colonization. BMC Biol 21, 98. 10.1186/s12915-023-01593-3.

13. Gross, A.C., Cam, H., Phelps, D.A., Saraf, A.J., Bid, H.K., Cam, M., London, C.A., Winget, S.A., Arnold, M.A., Brandolini, L., Mo, X., et al. (2018). IL-6 and CXCL8 mediate osteosarcoma-lung interactions critical to metastasis. JCI Insight 3. 10.1172/jci.insight.99791.

14. Reinecke, J.B., Gross, A.C., Cam, M., Garcia, L.J., Cannon, M.V., Dries, R., Gryder, B.E., and Roberts, R.D. (2024). Aberrant activation of wound healing programs within the metastatic niche facilitates lung colonization by osteosarcoma cells. bioRxiv, 2024.2001.2010.575008. 10.1101/2024.01.10.575008.

15. Riggi, N., Suvà, M.L., and Stamenkovic, I. (2021). Ewing’s Sarcoma. New England Journal of Medicine 384, 154–164. 10.1056/NEJMra2028910.

16. Friedman, G.K., Beierle, E.A., Gillespie, G.Y., Markert, J.M., Waters, A.M., Chen, C.-Y., Denton, N.L., Haworth, K.B., Hutzen, B., Leddon, J.L., Streby, K.A., et al. (2015). Pediatric cancer gone viral. Part II: potential clinical application of oncolytic herpes simplex virus-1 in children. Molecular Therapy - Oncolytics 2, 15016. 10.1038/mto.2015.16.

17. Streby, K.A., Geller, J.I., Currier, M.A., Warren, P.S., Racadio, J.M., Towbin, A.J., Vaughan, M.R., Triplet, M., Ott-Napier, K., Dishman, D.J., Backus, L.R., et al. (2017). Intratumoral Injection of HSV1716, an Oncolytic Herpes Virus, Is Safe and Shows Evidence of Immune Response and Viral Replication in Young Cancer Patients. Clin Cancer Res 23, 3566–3574. 10.1158/1078-0432.CCR-16-2900.

18. Streby, K.A., Currier, M.A., Triplet, M., Ott, K., Dishman, D.J., Vaughan, M.R., Ranalli, M.A., Setty, B., Skeens, M.A., Whiteside, S., Yeager, N.D., et al. (2019). First-in-Human Intravenous Seprehvir in Young Cancer Patients: A Phase 1 Clinical Trial. Mol Ther 27, 1930–1938. 10.1016/j.ymthe.2019.08.020.

19. Barone, A., Chi, D.-C., Theoret, M.R., Chen, H., He, K., Kufrin, D., Helms, W.S., Subramaniam, S., Zhao, H., Patel, A., Goldberg, K.B., et al. (2017). FDA Approval Summary: Trabectedin for Unresectable or Metastatic Liposarcoma or Leiomyosarcoma Following an Anthracycline-Containing Regimen. Clinical Cancer Research 23, 7448–7453. 10.1158/1078-0432.Ccr-17-0898.

20. Germano, G., Frapolli, R., Belgiovine, C., Anselmo, A., Pesce, S., Liguori, M., Erba, E., Uboldi, S., Zucchetti, M., Pasqualini, F., Nebuloni, M., et al. (2013). Role of macrophage targeting in the antitumor activity of trabectedin. Cancer Cell 23, 249–262. 10.1016/j.ccr.2013.01.008.

21. D’Incalci, M., Badri, N., Galmarini, C.M., and Allavena, P. (2014). Trabectedin, a drug acting on both cancer cells and the tumour microenvironment. Br J Cancer 111, 646–650. 10.1038/bjc.2014.149.

22. Banerjee, P., Zhang, R., Ivan, C., Galletti, G., Clise-Dwyer, K., Barbaglio, F., Scarfo, L., Aracil, M., Klein, C., Wierda, W., Plunkett, W., et al. (2019). Trabectedin Reveals a Strategy of Immunomodulation in Chronic Lymphocytic Leukemia. Cancer Immunol Res 7, 2036–2051. 10.1158/2326-6066.CIR-19-0152.

23. Belgiovine, C., Frapolli, R., Liguori, M., Digifico, E., Colombo, F.S., Meroni, M., Allavena, P., and D’Incalci, M. (2021). Inhibition of tumor-associated macrophages by trabectedin improves the antitumor adaptive immunity in response to anti-PD-1 therapy. Eur J Immunol 51, 2677–2686. 10.1002/eji.202149379.

24. Borgoni, S., Iannello, A., Cutrupi, S., Allavena, P., D’Incalci, M., Novelli, F., and Cappello, P. (2018). Depletion of tumor-associated macrophages switches the epigenetic profile of pancreatic cancer infiltrating T cells and restores their anti-tumor phenotype. Oncoimmunology 7, e1393596. 10.1080/2162402x.2017.1393596.

25. D’Incalci, M., and Galmarini, C.M. (2010). A review of trabectedin (ET-743): a unique mechanism of action. Mol Cancer Ther 9, 2157–2163. 10.1158/1535-7163.MCT-10-0263.

26. Grohar, P.J., Griffin, L.B., Yeung, C., Chen, Q.R., Pommier, Y., Khanna, C., Khan, J., and Helman, L.J. (2011). Ecteinascidin 743 interferes with the activity of EWS-FLI1 in Ewing sarcoma cells. Neoplasia 13, 145–153. 10.1593/neo.101202.

27. Grohar, P.J., Segars, L.E., Yeung, C., Pommier, Y., D’Incalci, M., Mendoza, A., and Helman, L.J. (2014). Dual targeting of EWS-FLI1 activity and the associated DNA damage response with trabectedin and SN38 synergistically inhibits Ewing sarcoma cell growth. Clin Cancer Res 20, 1190–1203. 10.1158/1078-0432.CCR-13-0901.

28. Harlow, M.L., Chasse, M.H., Boguslawski, E.A., Sorensen, K.M., Gedminas, J.M., Kitchen-Goosen, S.M., Rothbart, S.B., Taslim, C., Lessnick, S.L., Peck, A.S., Madaj, Z.B., et al. (2019). Trabectedin Inhibits EWS-FLI1 and Evicts SWI/SNF from Chromatin in a Schedule-dependent Manner. Clin Cancer Res 25, 3417–3429. 10.1158/1078-0432.CCR-18-3511.

29. Hernando-Cubero, J., Sanz-Moncasi, P., Hernández-García, A., Pajares-Bernard, I., and Martínez-Trufero, J. (2016). Metastatic extraskeletal Ewing’s sarcoma treated with trabectedin: A case report. Oncol Lett 12, 2936–2941. 10.3892/ol.2016.4950.

30. Herzog, J., von Klot-Heydenfeldt, F., Jabar, S., Ranft, A., Rossig, C., Dirksen, U., Van den Brande, J., D’Incalci, M., von Luettichau, I., Grohar, P.J., Berdel, W.E., et al. (2016). Trabectedin Followed by Irinotecan Can Stabilize Disease in Advanced Translocation-Positive Sarcomas with Acceptable Toxicity. Sarcoma 2016, 7461783. 10.1155/2016/7461783.

31. Lau, L., Supko, J.G., Blaney, S., Hershon, L., Seibel, N., Krailo, M., Qu, W., Malkin, D., Jimeno, J., Bernstein, M., and Baruchel, S. (2005). A phase I and pharmacokinetic study of ecteinascidin-743 (Yondelis) in children with refractory solid tumors. A Children’s Oncology Group study. Clin Cancer Res 11, 672–677.

32. Baruchel, S., Pappo, A., Krailo, M., Baker, K.S., Wu, B., Villaluna, D., Lee-Scott, M., Adamson, P.C., and Blaney, S.M. (2012). A phase 2 trial of trabectedin in children with recurrent rhabdomyosarcoma, Ewing sarcoma and non-rhabdomyosarcoma soft tissue sarcomas: a report from the Children’s Oncology Group. Eur J Cancer 48, 579–585. 10.1016/j.ejca.2011.09.027.

33. Grohar, P., Ballman, K.V., Heise, R., Glod, J., Wedekind, M.F., Mascarenhas, L., Gedminas, J.M., DuBois, S.G., Maki, R.G., Crompton, B.D., Hayashi, M., et al. (2023). SARC037: Results of phase I study of trabectedin given as a 1-hour (h) infusion in combination with low dose irinotecan in relapsed/refractory Ewing sarcoma (ES). Journal of Clinical Oncology 41, 11519–11519. 10.1200/JCO.2023.41.16_suppl.11519.

34. Denton, N.L., Chen, C.Y., Hutzen, B., Currier, M.A., Scott, T., Nartker, B., Leddon, J.L., Wang, P.Y., Srinivas, R., Cassady, K.A., Goins, W.F., et al. (2018). Myelolytic Treatments Enhance Oncolytic Herpes Virotherapy in Models of Ewing Sarcoma by Modulating the Immune Microenvironment. Mol Ther Oncolytics 11, 62–74. 10.1016/j.omto.2018.10.001.

35. Ghilas, S., Ambrosini, M., Cancel, J.C., Brousse, C., Massé, M., Lelouard, H., Dalod, M., and Crozat, K. (2021). Natural killer cells and dendritic epidermal γδ T cells orchestrate type 1 conventional DC spatiotemporal repositioning toward CD8(+) T cells. iScience 24, 103059. 10.1016/j.isci.2021.103059.

36. Kroczek, R., and Henn, V. (2012). The Role of XCR1 and its Ligand XCL1 in Antigen Cross-Presentation by Murine and Human Dendritic Cells. Frontiers in Immunology 3. 10.3389/fimmu.2012.00014.

37. Böttcher, J.P., Bonavita, E., Chakravarty, P., Blees, H., Cabeza-Cabrerizo, M., Sammicheli, S., Rogers, N.C., Sahai, E., Zelenay, S., and Reis e Sousa, C. (2018). NK Cells Stimulate Recruitment of cDC1 into the Tumor Microenvironment Promoting Cancer Immune Control. Cell 172, 1022–1037.e1014. 10.1016/j.cell.2018.01.004.

38. Kalamvoki, M., and Deschamps, T. (2016). Extracellular vesicles during Herpes Simplex Virus type 1 infection: an inquire. Virology Journal 13, 63. 10.1186/s12985-016-0518-2.

39. Kalamvoki, M., Du, T., and Roizman, B. (2014). Cells infected with herpes simplex virus 1 export to uninfected cells exosomes containing STING, viral mRNAs, and microRNAs. Proc Natl Acad Sci U S A 111, E4991–4996. 10.1073/pnas.1419338111.

40. 40. Cucè, M., Gallo Cantafio, M.E., Siciliano, M.A., Riillo, C., Caracciolo, D., Scionti, F., Staropoli, N., Zuccalà, V., Maltese, L., Di Vito, A., Grillone, K., et al. (2019). Trabectedin triggers direct and NK-mediated cytotoxicity in multiple myeloma. J Hematol Oncol 12, 32. 10.1186/s13045-019-0714-9.

41. Ratti, C., Botti, L., Cancila, V., Galvan, S., Torselli, I., Garofalo, C., Manara, M.C., Bongiovanni, L., Valenti, C.F., Burocchi, A., Parenza, M., et al. (2017). Trabectedin Overrides Osteosarcoma Differentiative Block and Reprograms the Tumor Immune Environment Enabling Effective Combination with Immune Checkpoint Inhibitors. Clin Cancer Res 23, 5149–5161. 10.1158/1078-0432.Ccr-16-3186.

42. Ramachandran, B., Rajkumar, T., and Gopisetty, G. (2021). Challenges in modeling EWS-FLI1-driven transgenic mouse model for Ewing sarcoma. Am J Transl Res 13, 12181–12194.

43. Mestas, J., and Hughes, C.C.W. (2004). Of Mice and Not Men: Differences between Mouse and Human Immunology. The Journal of Immunology 172, 2731–2738. 10.4049/jimmunol.172.5.2731.

44. Vasileva, E., Warren, M., Triche, T.J., and Amatruda, J.F. (2022). Dysregulated heparan sulfate proteoglycan metabolism promotes Ewing sarcoma tumor growth. Elife 11. 10.7554/eLife.69734.

45. Daley, J.D., Mukherjee, E.M., Cillo, A.R., Tufino, A.C., Bailey, N.G., Bruno, T.C., McAllister-Lucas, L.M., Vignali, D.A., and Bailey, K.M. (2022). Abstract A004: Radiation-induced changes to the immune microenvironment in an immunocompetent mouse model of Ewing sarcoma. Clinical Cancer Research 28, A004–A004. 10.1158/1557-3265.Sarcomas22-a004.

46. Takahashi, S., Araki, N., Sugiura, H., Ueda, T., Takahashi, M., Morioka, H., Yonemoto, T., Hiraga, H., Hiruma, T., Kunisada, T., Matsumine, A., et al. (2014). A randomized phase II study comparing trabectedin (T) and best supportive care (BSC) in patients (pts) with translocation-related sarcomas (TRS). Journal of Clinical Oncology 32, 10524–10524. 10.1200/jco.2014.32.15_suppl.10524.

47. Baruchel, S., Pappo, A., Krailo, M., Baker, K.S., Wu, B., Villaluna, D., Lee-Scott, M., Adamson, P.C., and Blaney, S.M. (2012). A phase 2 trial of trabectedin in children with recurrent rhabdomyosarcoma, Ewing sarcoma and non-rhabdomyosarcoma soft tissue sarcomas: A report from the Children’s Oncology Group. European Journal of Cancer 48, 579–585. 10.1016/j.ejca.2011.09.027.

48. Chawla, S.P., Tellez, W.A., Chomoyan, H., Valencia, C., Ahari, A., Omelchenko, N., Makrievski, S., Brigham, D.A., Chua-Alcala, V., Quon, D., Moradkhani, A., et al. (2023). Activity of TNT: a phase 2 study using talimogene laherparepvec, nivolumab and trabectedin for previously treated patients with advanced sarcomas (NCT# 03886311). Front Oncol 13, 1116937. 10.3389/fonc.2023.1116937.

49. Khanna, C., Prehn, J., Yeung, C., Caylor, J., Tsokos, M., and Helman, L. (2000). An orthotopic model of murine osteosarcoma with clonally related variants differing in pulmonary metastatic potential. Clinical & Experimental Metastasis 18, 261–271. 10.1023/A:1006767007547.

50. Zhao, S., Kurenbekova, L., Gao, Y., Roos, A., Creighton, C.J., Rao, P., Hicks, J., Man, T.K., Lau, C., Brown, A.M.C., Jones, S.N., et al. (2015). NKD2, a negative regulator of Wnt signaling, suppresses tumor growth and metastasis in osteosarcoma. Oncogene 34, 5069–5079. 10.1038/onc.2014.429.

51. Chase, M., Chung, R.Y., and Chiocca, E.A. (1998). An oncolytic viral mutant that delivers the CYP2B1 transgene and augments cyclophosphamide chemotherapy. Nat Biotechnol 16, 444–448. 10.1038/nbt0598-444.

52. Leddon, J.L., Chen, C.Y., Currier, M.A., Wang, P.Y., Jung, F.A., Denton, N.L., Cripe, K.M., Haworth, K.B., Arnold, M.A., Gross, A.C., Eubank, T.D., et al. (2015). Oncolytic HSV virotherapy in murine sarcomas differentially triggers an antitumor T-cell response in the absence of virus permissivity. Mol Ther Oncolytics 1, 14010. 10.1038/mto.2014.10.

53. Hao, Y., Hao, S., Andersen-Nissen, E., Mauck, W.M., Zheng, S., Butler, A., Lee, M.J., Wilk, A.J., Darby, C., Zager, M., Hoffman, P., et al. (2021). Integrated analysis of multimodal single-cell data. Cell 184, 3573–3587.e3529. 10.1016/j.cell.2021.04.048.

54. Wu, T., Hu, E., Xu, S., Chen, M., Guo, P., Dai, Z., Feng, T., Zhou, L., Tang, W., Zhan, L., Fu, X., et al. (2021). clusterProfiler 4.0: A universal enrichment tool for interpreting omics data. The Innovation 2, 100141. 10.1016/j.xinn.2021.100141.

55. Yu, G., Wang, L.-G., Han, Y., and He, Q.-Y. (2012). clusterProfiler: an R Package for Comparing Biological Themes Among Gene Clusters. OMICS: A Journal of Integrative Biology 16, 284–287. 10.1089/omi.2011.0118.

56. Benayoun, B.A., Pollina, E.A., Singh, P.P., Mahmoudi, S., Harel, I., Casey, K.M., Dulken, B.W., Kundaje, A., and Brunet, A. (2019). Remodeling of epigenome and transcriptome landscapes with aging in mice reveals widespread induction of inflammatory responses. Genome Res 29, 697–709. 10.1101/gr.240093.118.

57. Aran, D., Looney, A.P., Liu, L., Wu, E., Fong, V., Hsu, A., Chak, S., Naikawadi, R.P., Wolters, P.J., Abate, A.R., Butte, A.J., et al. (2019). Reference-based analysis of lung single-cell sequencing reveals a transitional profibrotic macrophage. Nat Immunol 20, 163–172. 10.1038/s41590-018-0276-y.

58. Heng, T.S.P., Painter, M.W., Elpek, K., Lukacs-Kornek, V., Mauermann, N., Turley, S.J., Koller, D., Kim, F.S., Wagers, A.J., Asinovski, N., Davis, S., et al. (2008). The Immunological Genome Project: networks of gene expression in immune cells. Nature Immunology 9, 1091–1094. 10.1038/ni1008-1091.

