## Supplemental Figures for "Trabectedin Enhances Oncolytic Virotherapy by Reducing Barriers to Virus Spread and Cytotoxic Immunity in Preclinical Pediatric Bone Sarcoma"

**Supplemental Methods**

*scRNAseq Treatment and Collection Determinations (Figure S3)*

A673 tumors were implanted in athymic nude mice and allowed to reach 300-400 $mm^3$ each (n = 1 per cohort). Mice were treated with trabectedin I.V. by tail vein injection on Day 0. The oHSV C134 was diluted in PBS to $1.0x10^7$ pfu/mL per mouse (each mouse receiving 100 µL) and administered via i.Tu. injection. The T1D3-treated mouse received oHSV on Day 0 and the tumor was harvested on Day 3 (T1D3: 1 oHSV treatment, collection 3 days after final treatment). The T2D1-treated mouse received oHSV on Day 0 and 2 and the tumor was harvested on Day 3 (T2D1: 2 oHSV treatments, collection 1 day after final treatment). The T2D3-treated mouse received oHSV on Day 0 and 2 and the tumor was harvested on Day 5 (T2D5: 2 oHSV treatments, collection 3 days after final treatment). One mouse remained untreated as an untreated control and the tumor was harvested on Day 3. Following tumor collection from treated mice and subsequent disassociation, microscopy of the cells stained with DAPI was used to determine the live cell counts for each scRNAseq preparation. Cellular infiltrate levels were determined from FACS analysis of each tumor collection.

*Toxicity Studies (Figure S5)*

In an attempt to combat the toxicity observed in tumor-bearing C57BL/6 mice yet maintain efficacy, further efficacy studies in the F420 tumor model included variation on trabectedin dosing. The treatment cohorts were tested with a single dose of trabectedin treatment I.V. (0.15 mg/kg) on Day 0 in F420 tumor-bearing B6-albino mice. Trabectedin dose de-escalation was also performed in F420 tumor-bearing B6-albino mice, using the regimen described above with the following treatment groups: HSV1716 (1.0 $x10^8$ pfu) + trabectedin (0.075 mg/kg,I.V.), HSV1716 (1.0 $x10^8$ pfu) + trabectedin (0.0375 mg/kg, I.V.), HSV1716 (1.0 $x10^8$ pfu) + trabectedin (0.0188 mg/kg, I.V.), HSV1716 (1.0 $x10^8$ pfu) + trabectedin (0.0094 mg/kg, I.V.), HSV1716 (1.0 $x10^8$ pfu) + trabectedin (0.0047 mg/kg, I.V.), HSV1716 (1.0 $x10^8$ pfu) + trabectedin (0.0023 mg/kg, I.V.). Additionally, fragmented trabectedin dosing was used to mitigate the observed toxicity. The same regimen of oHSV and PBS as in the original efficacy was utilized, but trabectedin was administered I.V. on Days 0, 2, 4, 6, 8 at 0.05 mg/kg in B6-albino mice. Finally, the addition of steroid treatment (dexamethasone) to each of the above cohorts aimed to lower the toxic effects observed with trabectedin in the C57BL/6 and B6-albino mice. Dexamethasone was administered via intraperitoneal (IP) injection at 40 mg/kg in vf = 250 mL (Days -1, -2, 5, and 6).

To assess the exacerbated toxic effect of trabectedin when combined with oHSV in C57BL/6 and B6-albino mice, F420 tumors (~200 $mm^3$) were established in B6-albino mice and treated as described in the efficacy studies, except with only one trabectedin treatment (Day 0). Animals were subjected to exsanguination with euthanasia being confirmed by cervical dislocation on Days 3 and 6 for cytokine analysis and Days 3 and 7 for serum chemistry for blood chemistry and tissue histology. Blood was collected in serum-separating tubes (Becton Dickinson, San Jose, CA) and allowed to clot at room temperature for 20 minutes. The clotted blood was then centrifuged at >2000xg for 20 minutes at room temperature. After collecting the supernatant (serum) and storing at −80°C, the frozen serum samples were shipped to IDEXX BioAnalytics (North Grafton, MA) for analysis on a Mouse Cytokine 25-Panel (62579) and a Standard Toxicology Panel (62794). Liver, lung, brain, and tumor were harvested for histopathology examination. Briefly, harvested tissues were fixed for 24 hours in 10% buffered formalin (Thermo Fisher Scientific), placed in tissue cassettes, and then transferred to 70% ethanol for 24 hours. Fixed tissues were paraffin-embedded by the Morphology Core at Nationwide Children's Hospital (Columbus, OH). Cutting of serial sections of tissues, hematoxylin and eosin staining, and histopathological evaluations by a licensed veterinary pathologist were performed by IDEXX BioAnalytics (North Grafton, MA). Further histological analysis of tissues was performed at Nationwide Children's Hospital (Columbus, OH). Briefly, unstained sections were stained for P-glycoprotein by the Nationwide Children's Hospital Morphology Core using the rabbit recombinant antibody MA5-32282 (Thermo Fisher Scientific) at 1:1000 dilution followed by Signal Boost IHC detection reagent (HRO Rabbit) from Cell Signaling (Danvers, MA) and then counterstained with Harris hematoxylin. Liver and tumor P- glycoprotein staining was then evaluated and scored (0, 1+, 2+, or 3+) by L.E.B.

**CHLA258**

| | PBS | Trab. | oHSV | oHSV + Trab. |
|---|---|---|---|---|
| **CR** | 0 | 3 | 0 | 7 |
| **PR** | 0 | 3 | 0 | 10 |
| **SD** | 0 | 0 | 0 | 2 |
| **PD** | 10 | 9 | 9 | 2 |

**EW5**

| | PBS | Trab. | oHSV | oHSV + Trab. |
|---|---|---|---|---|
| **PR** | 0 | 0 | 6 | 9 |
| **SD** | 0 | 1 | 9 | 7 |
| **PD** | 10 | 20 | 2 | 3 |

**PDX-0027**

| | PBS | Trab. | oHSV | oHSV + Trab. |
|---|---|---|---|---|
| **CR** | 0 | 0 | 1 | 6 |
| **PR** | 0 | 0 | 3 | 0 |
| **SD** | 0 | 0 | 1 | 0 |
| **PD** | 4 | 7 | 0 | 0 |

**A673**

| | PBS | Trab. | oHSV | oHSV+Trab. |
|---|---|---|---|---|
| **CR** | 0 | 0 | 0 | 1 |
| **PR** | 0 | 0 | 3 | 5 |
| **SD** | 0 | 0 | 2 | 4 |
| **PD** | 8 | 6 | 3 | 1 |

**A673 in NSG**

| | PBS | Trab. | oHSV | oHSV + Trab. |
|---|---|---|---|---|
| **SD** | 0 | 0 | 0 | 2 |
| **PD** | 7 | 11 | 8 | 7 |

**F420**

| | PBS | Trab. | oHSV | oHSV + Trab. |
|---|---|---|---|---|
| **CR** | 0 | 1 | 0 | 2 |
| **PR** | 0 | 1 | 2 | 2 |
| **SD** | 0 | 1 | 0 | 2 |
| **PD** | 4 | 6 | 8 | 0 |

**K7M2**

| | PBS | Trab. | oHSV | oHSV + Trab. |
|---|---|---|---|---|
| **CR** | 0 | 1 | 0 | 5 |
| **PR** | 0 | 1 | 0 | 3 |
| **SD** | 0 | 0 | 1 | 0 |
| **PD** | 4 | 7 | 4 | 0 |

**K7M2 in nude mice**

| | PBS | Trab. | oHSV | oHSV + Trab. |
|---|---|---|---|---|
| **PR** | 0 | 0 | 0 | 1 |
| **PD** | 5 | 6 | 5 | 7 |

**K7M2 with NK cell depletion**

| | PBS | oHSV + Trab. + control serum | oHSV + Trab. | oHSV + Trab. + anti-NK |
|---|---|---|---|---|
| **CR** | 0 | 2 | 2 | 2 |
| **PR** | 0 | 1 | 3 | 2 |
| **SD** | 0 | 1 | 0 | 0 |
| **PD** | 8 | 1 | 1 | 4 |

**K7M2 with T cell depletion**

| | oHSV + Trab. | oHSV + Trab. + anti-CD4 | oHSV + Trab. + anti-CD8 | oHSV + Trab. + anti-CD4 + anti-CD8 |
|---|---|---|---|---|
| **CR** | 3 | 3 | 0 | 0 |
| **PR** | 1 | 0 | 2 | 3 |
| **PD** | 1 | 3 | 5 | 3 |

**Figure S1. Quantification of best responses for all mouse efficacy studies.** CR, PR, SD, or PD were assigned and quantified for the 28 day treatment response period for all mouse efficacy studies. These best responses were used to determine disease stabilization rates and best overall response rates for each efficacy study.

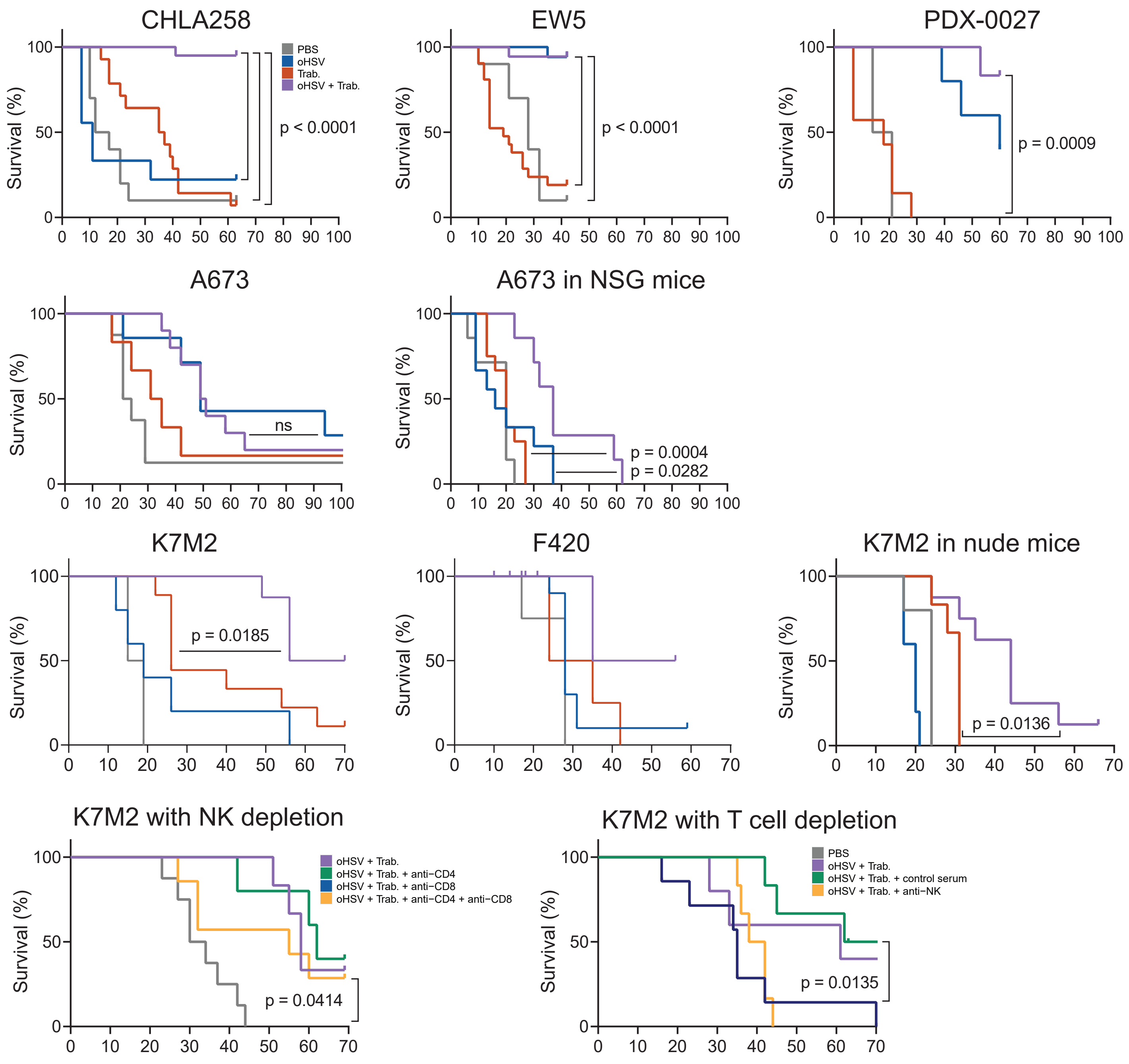

**Figure S2. Survival curves demonstrate the treatment-related survival advantages for all mouse models.** Survival curves were plotted for each mouse efficacy study. Statistical tests were performed using the Mantel-Cox log-rank test. Tick marks represent a survival event that is not a tumor burden-related death.

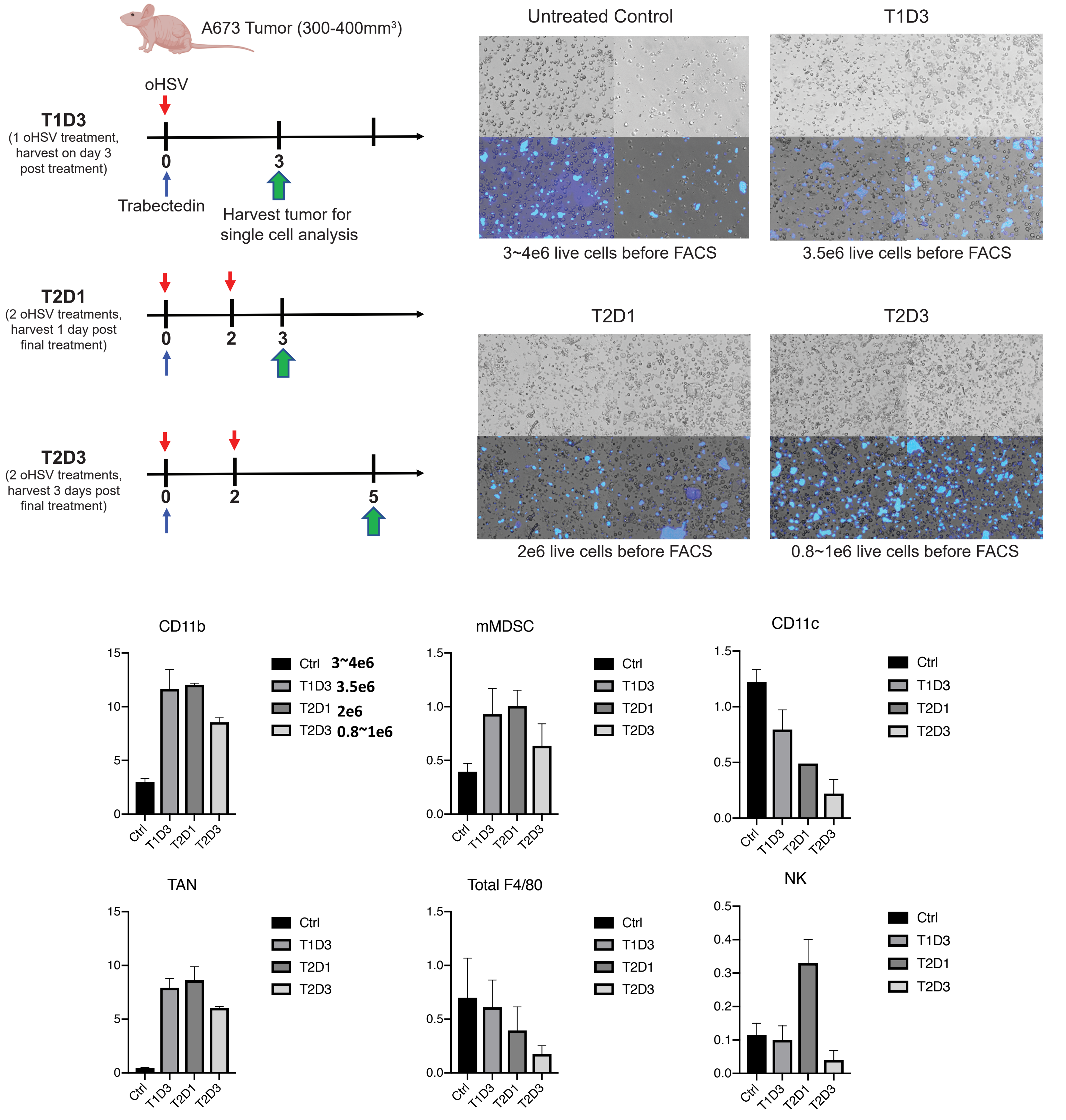

**Figure S3. Two oHSV treatments and a third day tumor harvest provided the best treatment regimen and collection time for optimal scRNAseq samples.** Numerous treatment regimens and collection times (shown in the schemata) were tested to determine the best scRNAseq sample quality. Following tumor collection from treated mice and subsequent disassociation, microscopy of the cells stained with DAPI was used to determine the live cell counts for each scRNAseq preparation. Cellular infiltrate levels were determined from FACS analysis of each tumor collection.

| | | avg_log2FC | pct.1 | pct.2 |
|---|---|---|---|---|
| | HSV1-RL2 | 0.01748651 | 0.026 | 0.007 |
| | HSV1-RS1 | 0.05501362 | 0.078 | 0.027 |
| Immediate Early Genes | HSV1-UL54 | 0.32190036 | 0.362 | 0.179 |
| | HSV1-US1 | 0.34115937 | 0.613 | 0.364 |
| | HSV1-US12 | 0.03539508 | 0.061 | 0.031 |
| | HSV1-UL23 | 0.1042827 | 0.153 | 0.094 |
| Early Genes | HSV1-UL29 | 0.2159139 | 0.182 | 0.084 |
| | HSV1-UL50 | 0.1822745 | 0.222 | 0.111 |
| | HSV1-UL2 | 0.1537922 | 0.167 | 0.083 |
| | HSV1-UL48 | 0.27210662 | 0.387 | 0.178 |
| | HSV1-UL19 | 0.08787415 | 0.081 | 0.045 |
| Late Genes | HSV1-UL27 | 0.22067906 | 0.293 | 0.15 |
| | HSV1-UL53 | 0.01433734 | 0.057 | 0.038 |
| | HSV1-UL44 | 0.04273984 | 0.053 | 0.029 |
| | HSV1-UL41 | 0.1050529 | 0.106 | 0.056 |

**Figure S4. Trabectedin increases the expression of HSV-1 genes associated with immediate early, early, and late infection time points.** The average fold change values (log2FC) for each time-dependent HSV-1 gene were calculated for oHSV+ Trabectedin- compared to oHSV-treated A673 tumors (Seurat). The percent of cells expressing each gene in tumor cells treated with oHSV+Trabectedin (pct.1) or oHSV alone (pct.2) are also shown.

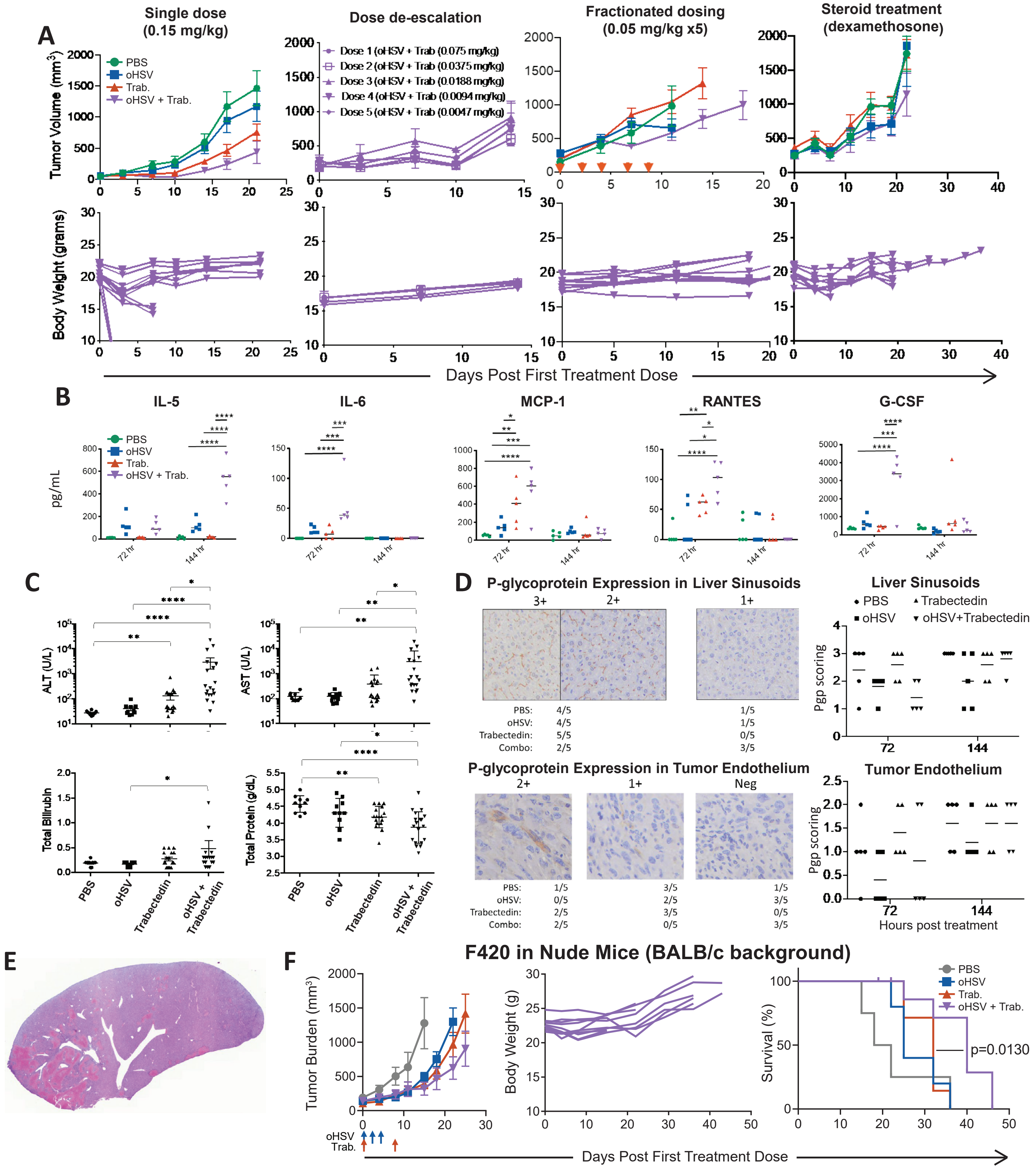

**Figure S5. Toxicity studies in the F420 tumor model reveal trabectedin-related, mouse model -dependent hepatotoxicity.** A) Trabectedin dose alteration–single dose, decreased concentration(s), fractionated dosing (0.05mg/kg on days 0, 2, 4, 6, 8), and steroid treatment (40mg/kg dexamethasone on days -2,-1, 5, 6)–resulted in body weight stabilization but higher tumor burdens. B) Cytokine and C) toxicology levels were quantified from blood samples of treated B6-albino mice. D) Liver and tumor P-glycoprotein histological expression and scoring was determined for treated tumor-bearing B6-albino mice. E) A liver histological section from a combination-treated B6-albino mouse displayed hepatotoxicity, with necrosis shown in red. F) Tumor burden, body weight, and survival curves for treated F420 Athymic nude mice (BALB/c background) showed safety but decreased combinatorial efficacy.

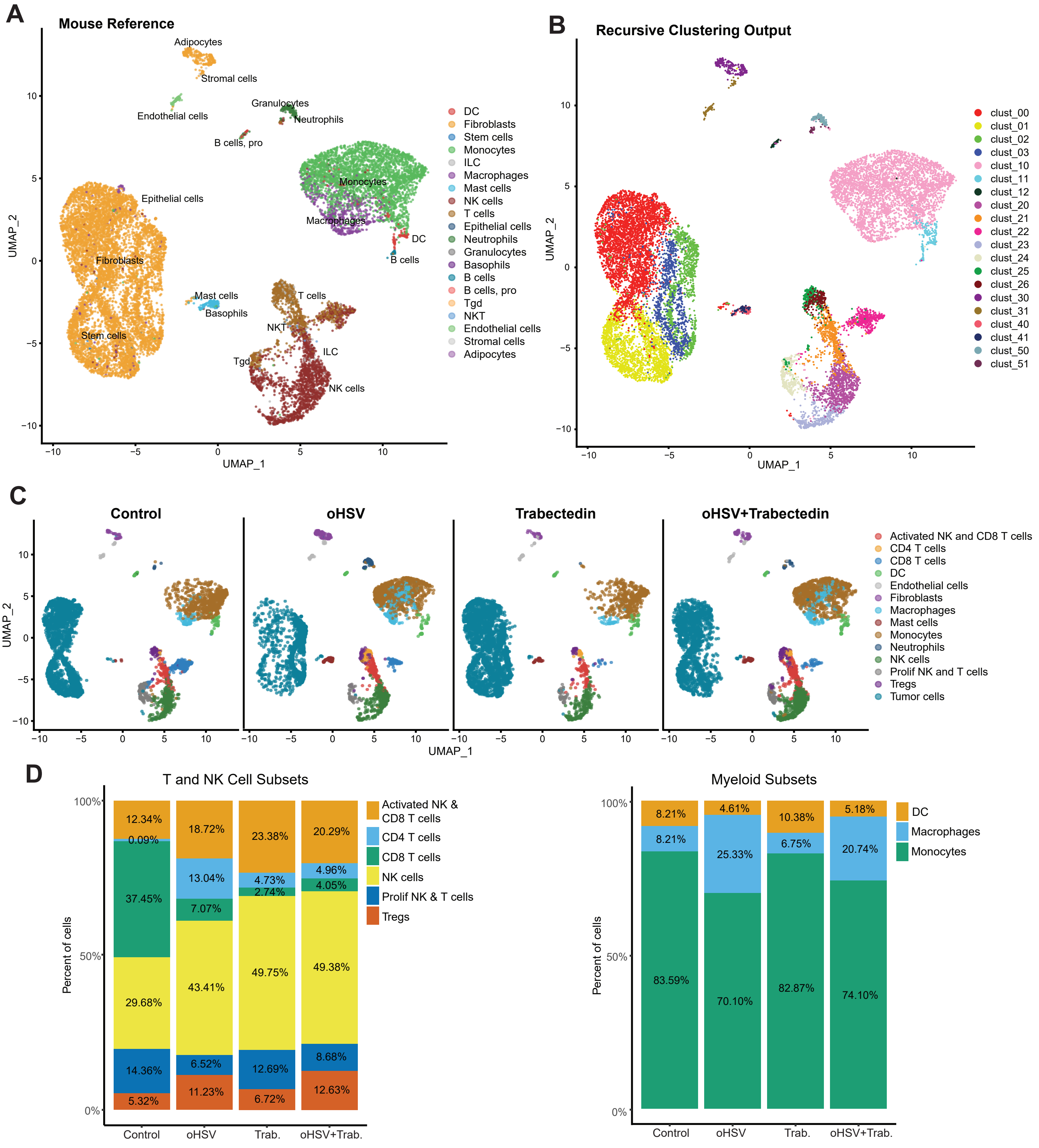

**Figure S6. scRNAseq analysis depicts the innate and adaptive immune cells infiltrating the osteosarcoma microenvironment.** A) The reference-based SingleR package annotated the dataset with mouse cell type assignment. B) The output from unsupervised recursive clustering shows distinct clusters related to cell type. C) Using these results and cell type-specific differential gene expression analysis, we assigned cell types and visualized these across treatment groups. D) Stacked barplots (ggplot2) demonstrated the relative proportion of specific cell types from two larger cohorts: cytotoxic lymphocytes and myeloid cells. The datasets for each treatment group were merged without a need for integration and plotted in the same UMAP space (Seurat).

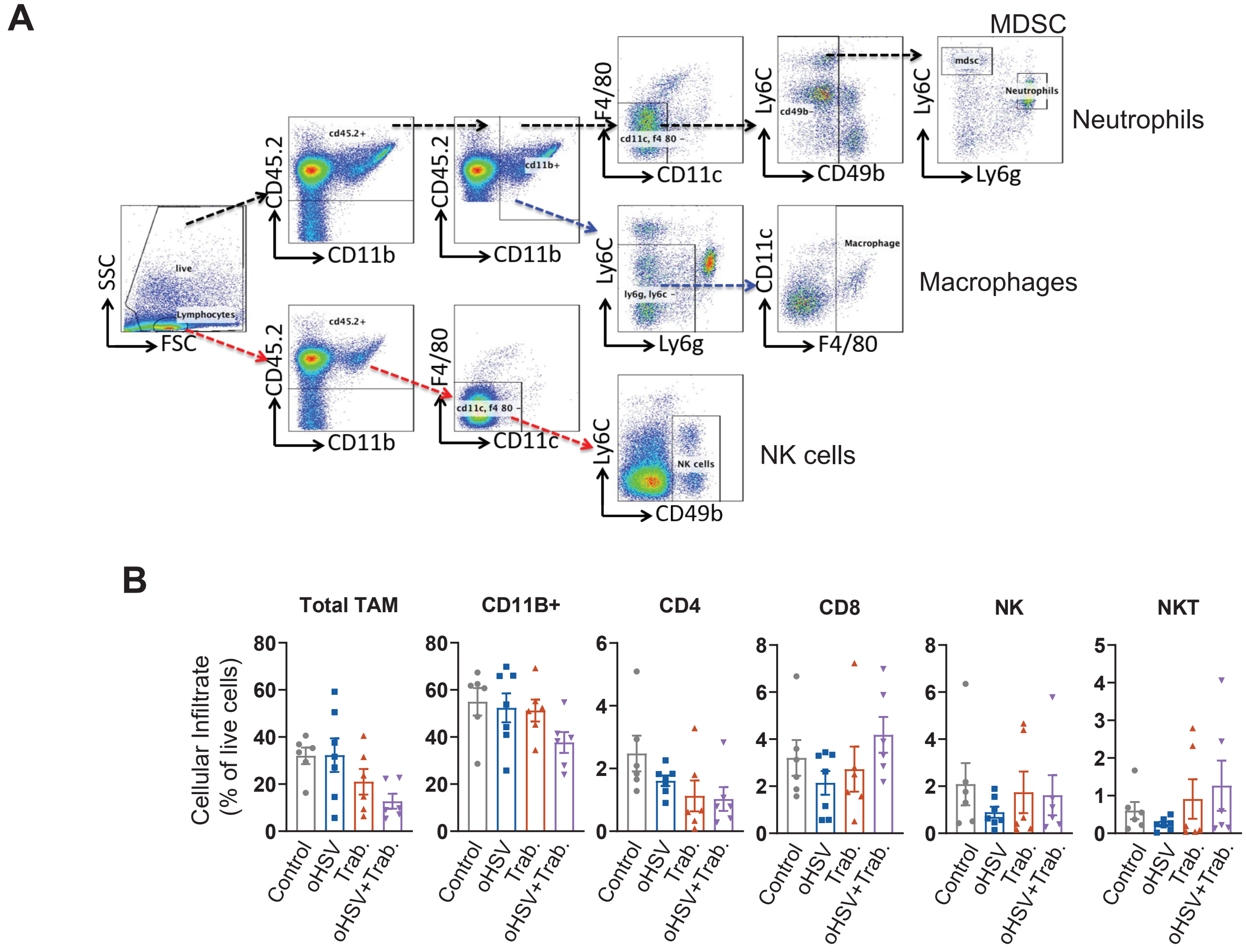

**Figure S7. FACS gating strategy and cellular infiltrates in treated-K7M2 tumors.** A) An example FACS gating strategy is shown for innate immune subsets. B) The FACS cellular infiltrate results that did not meet significance ($p<0.05$) from the treated-K7M2 tumors demonstrate minor changes in cellular infiltrates after treatment. The treatment regimen follows the previously described methodology for Figure 4.
